# Loss of heterochromatin and retrotransposon silencing constitute an early phase in oocyte aging

**DOI:** 10.1101/2020.10.28.358440

**Authors:** Peera Wasserzug-Pash, Rachel Rothman, Eli Reich, Oshrat Schonberger, Yifat Weiss, Naama Srebnik, Yaara Cohen-Hadad, Amir Weintraub, Ido Ben-Ami, Hananel Holzer, Michael Klutstein

## Abstract

Mammalian oocyte quality reduces with female age. A well-studied aspect of this deterioration is an age-associated rise in oocyte aneuploidy. We show that prior to the occurrence of significant aneuploidy (at the age of 9 months in mouse females), epigenetic changes occur and impact oocyte quality and maturation ability. At this age- we observe a reduction in heterochromatin marks in mouse oocytes. This decrease is apparent in both constitutive heterochromatin and facultative heterochromatin marks but is absent in active euchromatic marks which remain constant. A decrease of heterochromatin marks with age is also observed in human GV oocytes from IVF treatments. Heterochromatin loss with age is associated with an elevation in retrotransposon RNA transcription and processing, an elevation in retrotransposon protein expression, elevation in DNA repair proteins nuclear localization and oocyte maturation defects. Artificial inhibition of the heterochromatin machinery in young oocytes causes an elevation in retrotransposon expression and processing and oocyte maturation defects. Collectively, our work demonstrates an early stage of oocyte aging, characterized by the loss of heterochromatin associated chromatin marks and activation of retrotransposons which cause DNA damage and impair oocyte maturation. We hypothesize that this heterochromatin loss serves as an oocyte associated “epigenetic clock” and is exploited by the cell as an oocyte QC mechanism.

**One Sentence Summary:** Oocyte aging includes an early pre-aneuploidy phase when loss of repressive chromatin marks occurs as well as retrotransposon activation and egg maturation defects.

## Introduction

Reproductive aging is defined as the age-related loss of fertility due to increasing damage to the reproductive and other systems. Oocytes themselves accumulate damage in an age-related manner (Igarashi et al., 2015) (Miao et al., 2009) and deteriorate to the point where they are non-fertile. In human females this occurs at a relatively early age, before the onset of aging in other organs and tissues. In our era of increased rate of delayed childbearing, it is becoming crucial to understand the mechanisms underlying the compromised quality of oocytes with age. Previous studies have demonstrated that oocytes have increased rates of aneuploidy with age (Hassold & Hunt, 2001) (Nagaoka et al., 2012). One of the causes for this is due to precautious sister chromatid separation and resulting non-disjunction (Gruhn et al., 2019) . This happens because of loss of sister chromatid cohesion due to the unbinding and inability to re-load Cohesin complexes containing Rec8 on chromosomes (Burkhardt et al., 2016) (Lister et al., 2010) (Gruhn et al., 2019).

Despite the dominance of aneuploidy as a contributing factor to oocyte aging and loss, it is clear that additional factors contribute to the process. The decrease in mature oocyte numbers and in female fertility occurs earlier than the onset of aneuploidy (Merriman et al., 2012), a fact that demonstrates the effect of these additional factors.

Changes in epigenetic regulation of gene expression and altered chromosome dynamics have been recognized as contributors to aging, and epigenetic changes during aging have been listed among the “hallmarks of aging” (Lopez-Otin et al., 2013) . The loss of heterochromatin histone marks has been associated with the aging process in many systems and tissues (Zhang et al., 2015) (Djeghloul et al., 2016) (Jeon et al., 2018) (Keenan et al., 2020). The consequences of heterochromatin de-regulation in aging may be related to the activated transcription of transposable elements (TE) in the genome, and their subsequent effect on genome stability and cellular integrity. This has been shown to occur in several organisms and systems (De Cecco et al., 2013) (Chen et al., 2016) (Patterson et al., 2015) (Dennis et al., 2012) (Tarallo et al., 2012).

It was shown that epigenetic changes occur in oocytes of advanced maternal age (Manosalva & Gonzalez, 2010) (Yue et al., 2012), but this was always shown at an age where aneuploidy has already occurred at high rates and therefore deciphering the exact causal relationship between infertility, aneuploidy and epigenetic changes has not been possible. It is also unclear whether TE are activated in older oocytes, and whether this phenomenon is related to oocyte aging phenotypes.

Here we show that epigenetic changes in oocytes can be detected at an age of 9 months in mice, an age reported to still have low levels of oocyte aneuploidy. We show that these changes are characterized by the loss of repressive histone marks, elevation of retrotransposon mRNA transcription, elevated processing of repeated sequences and retrotransposons and increased activation of the DNA repair machinery. Treatment of oocytes with epigenetic drugs that inhibit heterochromatin formation can mimic the effect of aging and cause a decrease in oocyte maturation rates and elevation in retrotransposon activity and DNA damage.

## Results

In order to investigate the epigenetic status of oocytes at an age before the onset of aneuploidy, we compared the levels of epigenetic markers by immunofluorescence (IF) in GV oocytes of normally ovulating (i.e. not super-ovulated) 2 months old (young) and 9 months old (old) mouse females. According to previous studies (Manosalva & Gonzalez, 2010) (Koehler et al., 2006), mouse oocyte aneuploidy at this age is low and not significantly different than in young mice, but the level of oocyte maturation at this age by IVM (in-vitro maturation), number of oocytes found in the oviduct after superovulation and fecundity is already markedly reduced. Despite the absence of significant levels of aneuploidy, the levels of the constitutive heterochromatin markers H3K9me2 and HP1γ decrease significantly with age when assessed by in-situ immuno-fluorescent staining (Fig. 1A, B and S1 see supplementary methods for further details). Interestingly, the decrease in heterochromatin signal is much more pronounced as the oocyte proceeds to the stage of surrounded nucleolus (SN)- when the oocyte undergoes transcriptional shut-down (Fig. S2). These results hint to the possibility of an increasing epigenetic defect during the transition between NSN and SN (which is accompanied by transcriptional silencing of the egg, (Kageyama et al., 2007) (Bonnet-Garnier et al., 2012)), at 9 months old eggs. In addition, the level of the facultative heterochromatin mark H3K27me3 also reduces with age when assessed by in-situ immuno-fluorescent staining (Fig. 1C, S1). This decrease occurs uniformly in all GV stage oocytes (both NSN and SN), showing that some age-related epigenetic defects can already be seen at an early oocyte developmental stage (Fig. S2).

**Figure 1:**
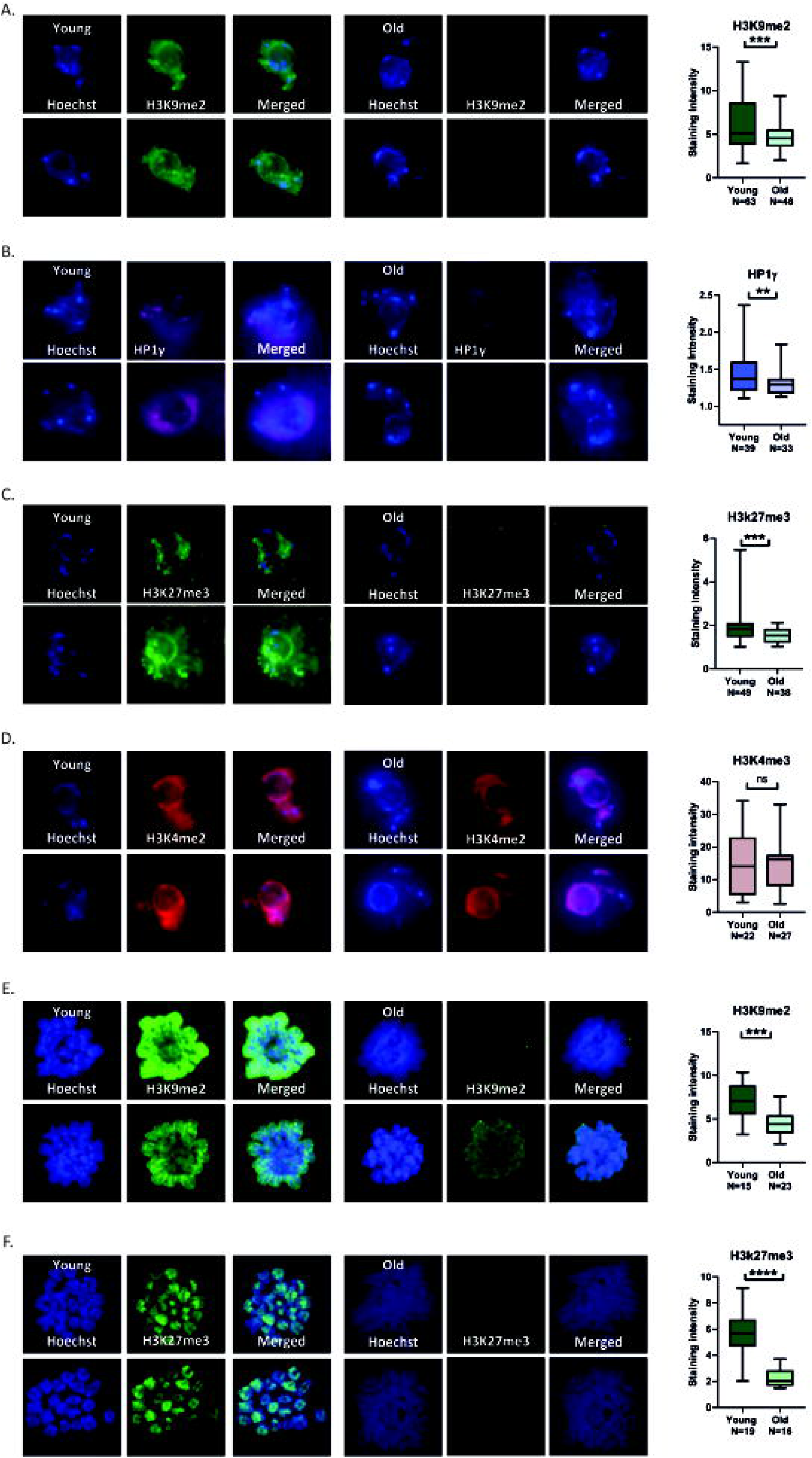
9 months old oocytes lose heterochromatin marks: (A-B) In situ staining of old and young GV mice oocytes, and staining intensity analysis, for constitutive heterochromatin markers (H3K9me2 and HP1γ) (C) In situ staining of old and young GV mice oocytes, and staining intensity analysis, for the facultative heterochromatin marker H3K27me3 (D) In situ staining of old and young GV mice oocytes, and staining intensity analysis, for the euchromatin marker H3K4me3 (E-F) staining of MI metaphase chromosomes (see Methods) of old and young oocytes chromosomes, for heterochromatin markers (H3k9me2, H3k27me3), and staining intensity analysis.

To investigate whether the epigenetic decrease in repressive marks occurs as a result of histone loss, and to control for differences in oocyte staining capacity between young and old oocytes, we stained oocytes by in-situ immuno-fluorescence for the active chromatin marks H3K27Ac and H3K4me3. Importantly, no significant difference was detected between young and old oocytes for these two chromatin marks (Fig. 1D, S3). These results show that the loss of repressive marks is specific to heterochromatin and that no detectable histone loss has occurred in old oocytes.

We also explored if the epigenetic defect persists in later stages of oocyte development. To do this- we explored the epigenome of oocytes at metaphase of MI by performing chromosome spreads followed by immuno-staining. Spreads were stained for H3K9me2, and for H3K27me3 (Fig. 1E, F, S1). In these experiments a significant drop in signal was observed in older oocytes. This result shows that the epigenetic defects that occur in older oocytes persist till the meiotic divisions. Since the spread experiments enable the visualization of the entire chromosome, we asked whether the decrease in repressive chromatin marks occurs on specific loci in the genome, or whether the decrease is uniform along the entire chromosome. Our results show that the decrease appears widespread and includes all parts of the nucleus and chromosomes without preference for a specific nuclear compartment or chromosome locus (Fig. S4).

To investigate the consequences of the epigenome de-regulation on the transcription of TE in oocytes, we examined RNA expression from two retrotransposon families: long interspersed nuclear element-1 (L1) and intracisternal A particle (IAP) which were shown to be expressed in the germline (Crichton et al., 2014) (Dupressoir & Heidmann, 1996) (Trelogan & Martin, 1995). Oocytes from older females had a roughly 2-fold increased expression of both L1 and IAP transcripts compared to young oocytes as assessed by quantitative RT-PCR (Fig 2A). We confirmed these findings by immunostaining oocytes for a protein of the L1 retrotransposon. L1-encoded ORF1p was detectable at low levels in young oocytes and increased in intensity in older oocytes (Fig. 2B, S5). In accordance with increased activity of TE, older oocytes show increased recruitment of DNA repair machinery indicative of DNA damage. This was shown by elevated Rad51 nuclear localization in older oocytes, and higher presence of γH2Ax loci (Gasior et al., 2006) (Fig. 2C,D, S5). Foci of γH2Ax are considered a bona fide marker of DNA damage and repair (Fernandez-Capetillo et al., 2004) . An elevated nuclear localization of Rad51 is also a marker for DNA damage, and was observed in HeLa and Hct116 cells after treatment with ionizing radiation (Gildemeister et al., 2009) .

**Figure 2:**
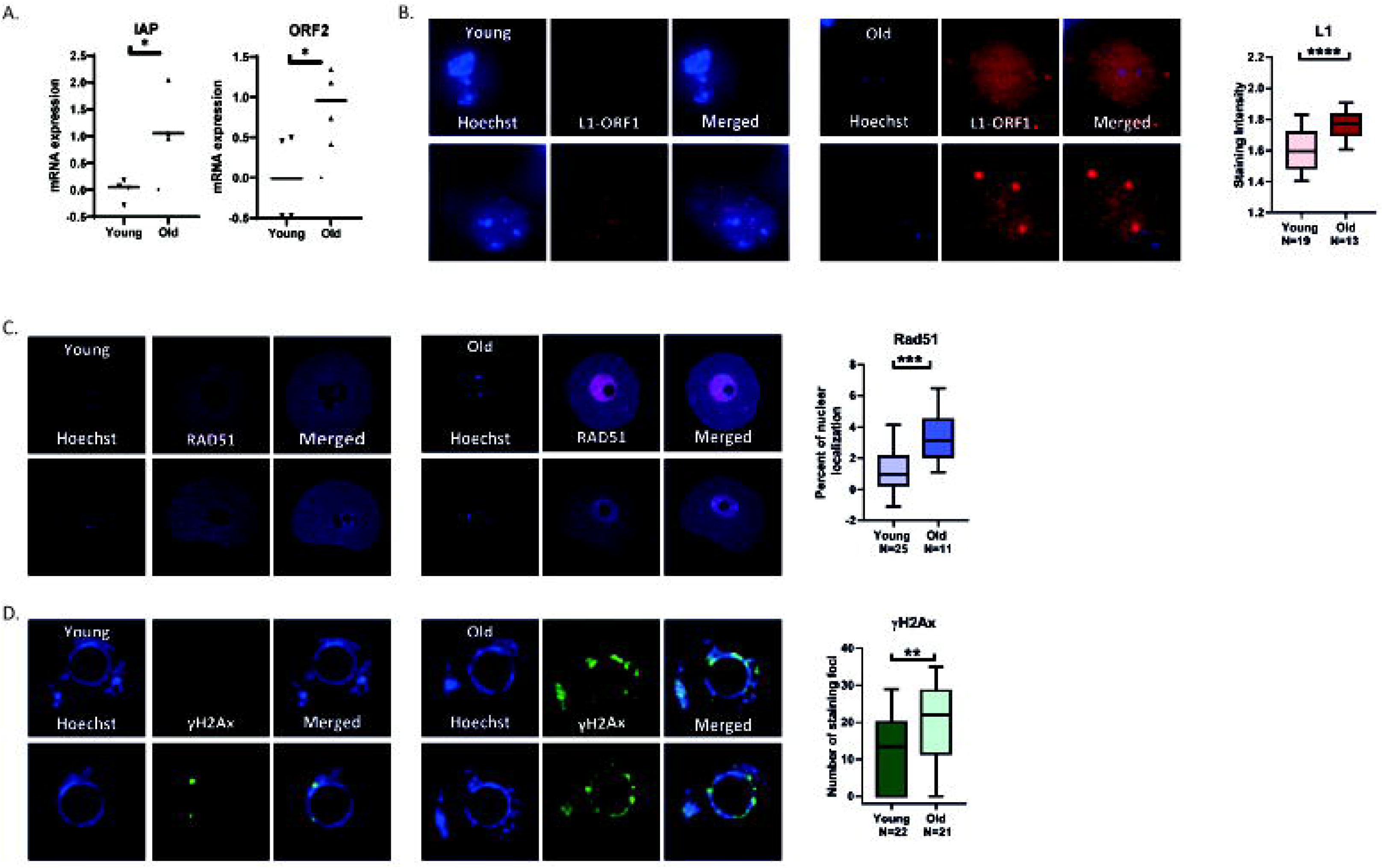
Old oocytes show elevated retrotransposon activation and DNA damage: (A) qRT-PCR measurement of L1 retrotransposon and IAP elements mRNA (see Methods) in young and old GV oocytes (B) In situ staining of L1- ORF1p and staining quantification in old and young GV mouse oocytes (C) In situ staining of Rad51 and quantification of nuclear localization in old and young GV mouse oocytes (D) In situ staining of γH2AX and quantification of number of nuclear foci in old and young GV mouse oocytes.

In addition to expression of retrotransposons in older oocyte, we investigated whether we also see evidence for increased processing of retrotransposon RNA. We therefore performed small RNA sequencing on oocytes from young and old females. We sequenced RNA molecules from 17 to 167 bp (median of 20 to 80 percentiles, see Methods and Fig. S6) with a median molecule size of 18 bp for the young oocytes and 106 for the older oocytes (Fig. S6A). a prominent peak of ribosomal RNA at 155 bp can be observed in both samples. The difference in small RNA sizes between young and old oocytes shows an elevated presence of RNA fragments of small sizes that not yet processed by Dicer, and could originate from spurious transcription due to loss of genomic repression of heterochromatin. The difference in RNA expression between the young and old oocytes was evident in the group of small RNAs coming from genomic repeats (and are usually marked by heterochromatin). Focusing on RNA from genomic repeats, we see that most repeat types remained unchanged between young and older oocytes (83%). The same amount of repeat types was overexpressed and under-expressed in older oocytes (9% and 8% respectively) (Fig. 3A and Table S1). The list of overexpressed repeat types in old oocytes (Fig. 3B, S6C, D and supplementary methods) shows that the predominant types of repeats that were overexpressed are LTR and IAP retrotransposon RNA molecules, while in the list of under-expressed repeat types also includes simple repeats and centromeric repeat transcripts. When looking specifically at retrotransposon repeats (386 out of 960 repeat types) the log2 fold change is increased compared to all repeat types (average 0.06 compared to 0.03) (Table S1 and Fig. S6E). In agreement with more retrotransposon mRNA and more retrotransposon-derived small RNA from some of the repeats in older oocytes, we also see a significant elevation in the signal of dsRNA molecules in older oocytes (Fig. 3C and Fig. S7). Moreover, staining for the Dicer protein, an enzyme which participates in the processing of retrotransposons (Soifer et al., 2005) (Svobodova et al., 2016)shows similar results (Fig. 3D and Fig. S7). We therefore conclude that the loss of epigenetic silencing in old eggs results in enhanced transcriptional activity and processing of retrotransposons in the genome. As previous studies (Flemr et al., 2013) (Tharp et al., 2020) have shown that elevated retrotransposon activity is detrimental to oocytes, we conclude that epigenetic aging damages oocytes partly through the activation of retrotransposons.

**Figure 3:**
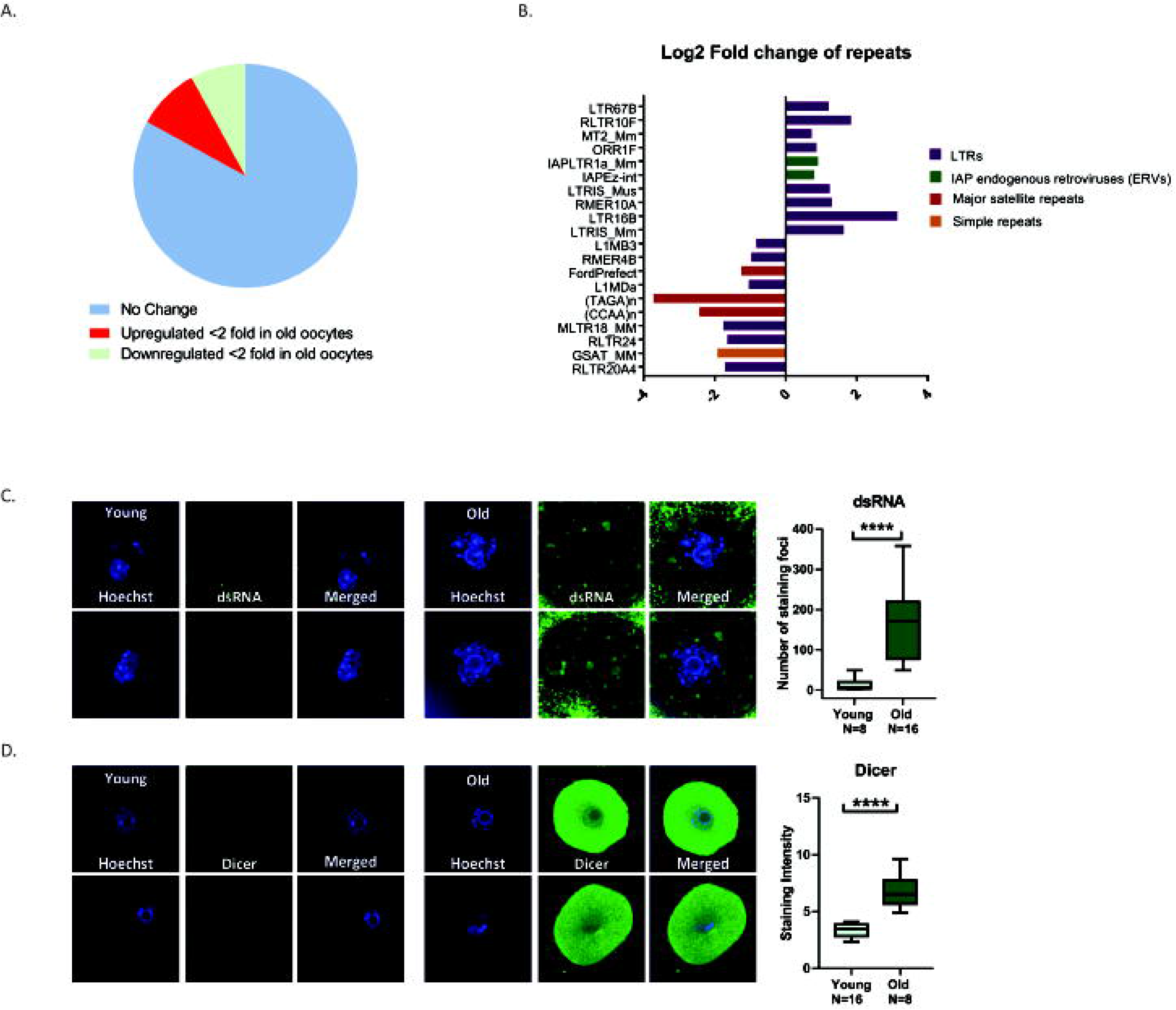
Old oocytes show elevated retrotransposon RNA processing activity: (A) Percentage of over-expressed and under-expressed types of repeat small RNA in old oocytes (B) Types of small RNAs from repeats that are over-expressed or under-expressed in old oocytes note dominance of LTR and IAP in over-expressed repeat types. (C) In situ staining for dsRNA and staining quantification in old and young GV mouse oocytes. (D) In situ staining for Dicer and staining quantification in old and young GV mouse oocytes. Note the elevation in signal for dsRNA and Dicer in old oocytes.

Since some significant differences in epigenetic regulation exist between mice and human (Hanna et al., 2018), we wanted to investigate whether heterochromatin loss with maternal age also occurs in human oocytes. For this purpose, we investigated the epigenome of GV oocytes from IVF treatments. As a general rule, GV oocytes are not used by clinics for fertilization, but to make sure the oocytes we used for research could not mature in-vitro, we waited another 24 hours with the oocytes in medium, before we fixed the immature GV oocytes and immune-stained them. Therefore, these oocytes likely represent a subset of human oocytes naturally arrested at the GV stage (for details of ethical approval see supplementary methods). Indeed, fixed oocytes that were stained for DNA visualization were arrested in several developmental stages (Fig. 4, S8). Further, we stained oocytes by in-situ immuno-fluorescent staining for H3K9me2. Stringent QC criteria were applied to the staining results in order to exclude eggs with fragmented or damaged genomes (see Methods). We found that in oocytes arrested both at the GV stage and after GVBD, there was an age-dependent decrease in H3K9me2 signal fitting a linear decreasing curve (Fig. 4, S8). As control, we also stained human oocytes for the Rec8 meiotic cohesin. This staining also shows (Fig. S8) a decreasing, age-dependent trend, as has been shown before for mouse and human cohesin (Liu & Keefe, 2008) (Chiang et al., 2010) (Tsutsumi et al., 2014).

**Figure 4:**
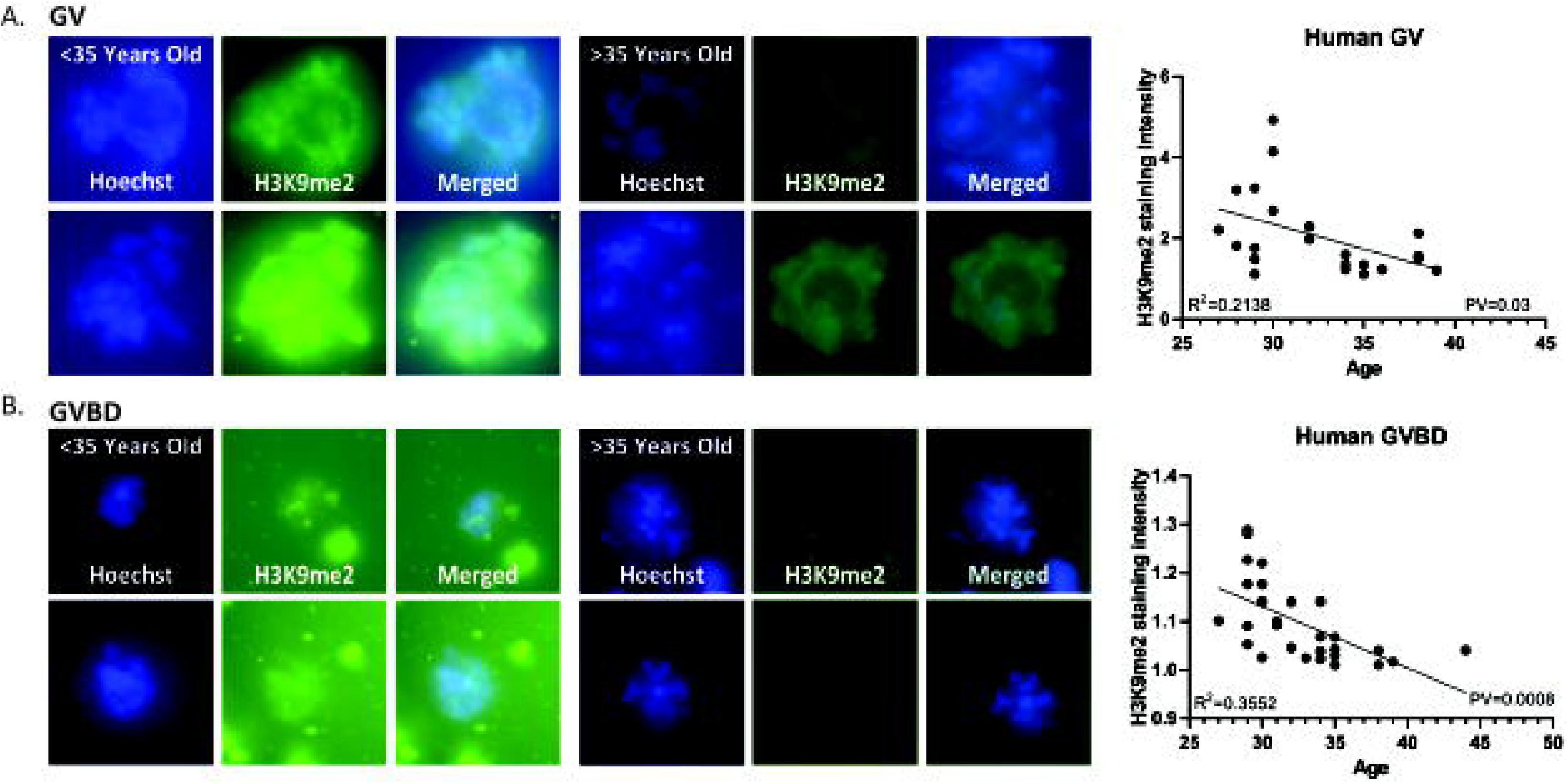
Human oocytes show a reduction of heterochromatin with age: A) In situ staining of GV human oocytes (N=21) for H3K9me2. These oocytes show a reduction of signal intensity with age in a linear regression curve (p=0.03). (B) (A) In situ staining of GVBD human oocytes (N=28) for H3K9me2. These oocytes show a reduction of signal intensity with age in a linear regression curve (p=0.0008).

In order to show a causal link between epigenetic deterioration and maturation defects of old oocytes we treated young mouse oocytes with chemicals known to affect epigenetic regulation. 2 months old oocytes were treated with the specific Suvar39h1/2 inhibitor Chaetocin (Greiner et al., 2005) at 0.5μM in vitro for 18h (Bertoldo et al., 2020). Staining for H3K9me2 decreased after the treatment by staining of chromosome spreads at metaphase of MI (Fig. S9). Assessment of the maturation efficiency of the oocytes after treatment with Chaeotocin shows a significant proportion of the oocytes did not properly mature after treatment (Fig. 5A). Staining of treated oocytes by in-situ immuno-fluorescent for the L1-encoded ORF1p shows an increase in staining intensity, as shown for naturally aged oocytes (Fig. 5B, S9). Consistently, dsRNA staining presented a significant increase in the treated compared to the untreated control oocytes (Fig. 5C, S9). Since heterochromatin that is characterized by H3K9me2 and binding of HP1γ is also associated with a histone de-acetylation on H3K27 (Naruse et al., 2020) (Wang et al., 2019), we sought to investigate whether inhibiting the deacetylation of histones in oocytes will cause a similar effect. Previous reports also showed that treatment with a histone deacetylase (HDAC) inhibitor affects oocyte maturation in-vitro (Jin et al., 2014). We thus treated young oocytes with the HDAC inhibitor Trichostatin A (TSA, 100nm for 4 h of arrest and then for 18h until MII) (Yoshida et al., 2003) In-vitro treated oocytes show an increase in H3K27Ac and also a decrease in H3K9me2 by in-situ staining linking the various heterochromatin features pathways such as H3K27 de-acetylation and H3K9 methylation in oocytes (Fig. S10). TSA treated young oocytes show similar maturation defects to Chaetocin treated young oocytes, and a marked elevation in dsRNA presence. Interestingly, TSA treated oocytes do not show a L1-encoded ORF1p elevation in signal. Instead L1-ORF1p accumulates in nuclei of treated oocytes, perhaps showing that de-acetylation activity is central to specific stages in retrotransposon maturation processes (Fig. S10), causing enhanced nuclear recruitment of the L1 protein when de-acetylation processes fail to occur. However, DNA damage response is elevated in TSA treated oocytes as shown by the elevated Rad51 nuclear localization in treated oocytes (Fig. S10). Collectively, these results show that epigenetic manipulation, in H3K9 methylation or H3K27 acetylation pathways causes the inability of young oocytes to mature, and mimics natural aging.

**Figure 5:**
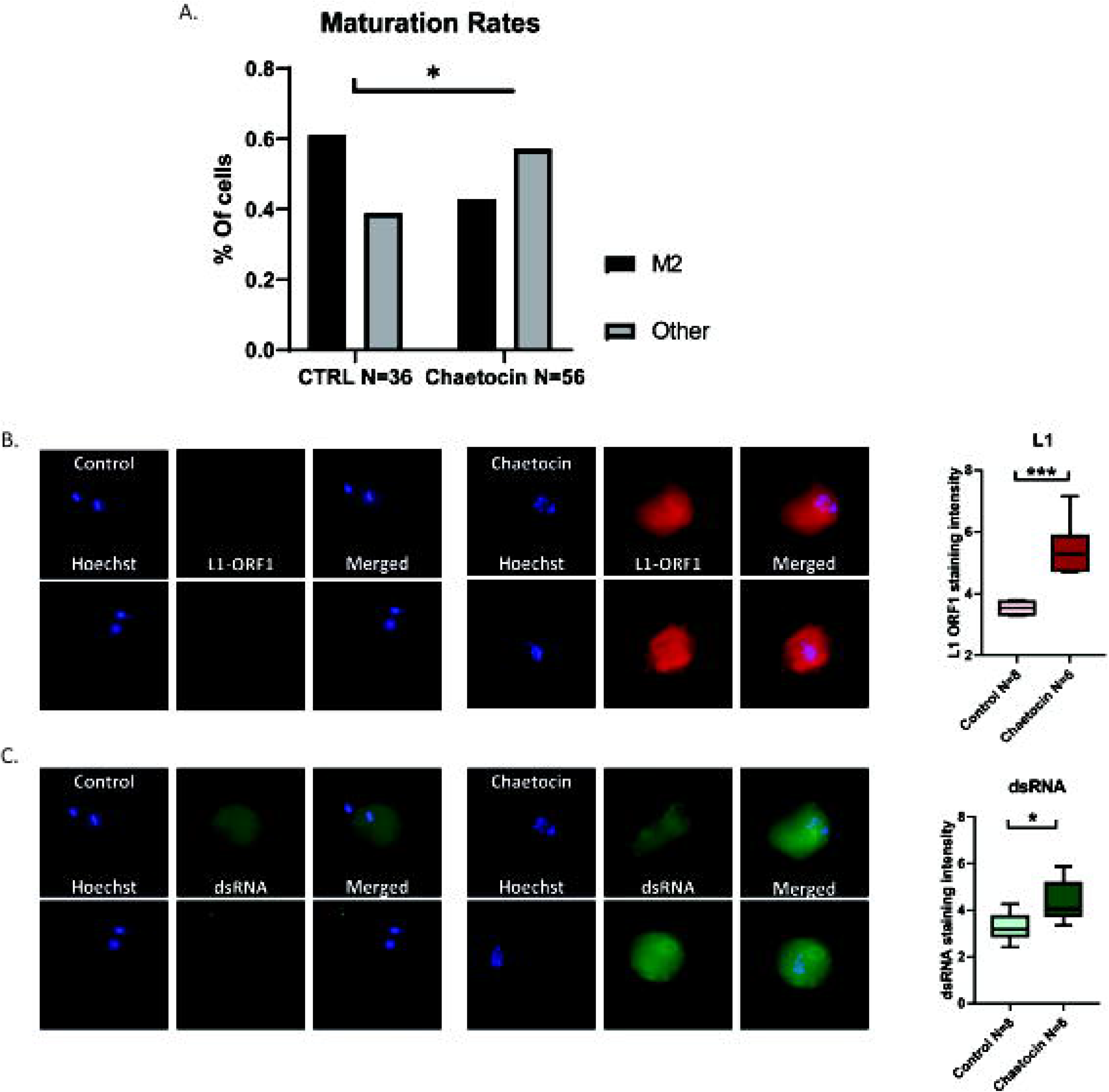
Treatment of oocytes with Chaetocin mimics natural aging: (A) Maturation efficiency of young GV mouse oocytes treated with Chaetocin and matured in-vitro (see Methods) (B) In situ staining for L1ORF1p and staining quantification in young GV mouse oocytes treated with Chaetocin. (C) In situ staining for dsRNA and staining quantification in young GV mouse oocytes treated with Chaetocin.

## Discussion

We show a reduction in repressive chromatin marks with age in mammalian oocytes. This trend can be observed in both mouse and human oocytes. In mouse oocytes, we show that heterochromatin loss occurs at a significant rate even before the onset of aneuploidy in oocytes. In a series of experiments, we demonstrate here that the reduction in repressive marks is associated with oocyte maturation defects, an increase in dsRNA and Dicer presence in the cytoplasm, an elevation in retrotransposon sRNA and an elevation in recruitment of DNA repair proteins. Using the epigenetic drugs Chaetocin and TSA we show that an artificial reduction in repressive marks in young oocytes also causes maturation defects and elevation in DNA damage. Our working model (Fig. S11) is therefore that due to external damage and erosion with time, epigenetic information and modified histones are lost from chromatin and replaced by non-modified nucleosomes. This leads to a cascade of molecular events that eventually lead to a re-structuring of the whole genome. Silencing of wide genomic domains is eventually lost, and retrotransposon RNA is transcribed. This in turn causes an elevation in DNA damage, and a decrease in oocyte maturation ability. This elevation in retrotransposon activity with age in oocytes could have survived in evolution in order to serve as a selection mechanism to eliminate non-functional oocytes from the oocyte pool. A similar mechanism has been previously reported to occur during fetal oocyte attrition (Tharp et al., 2020) (Malki et al., 2014). We thus hypothesize that heterochromatin loss serves as an oocyte associated ‘epigenetic clock” in a similar way to processes which occur (at a later age) in other cell types and is exploited by the cell as an oocyte QC mechanism.

---

**Figure S1: 9 months old oocytes lose heterochromatin marks (additional examples):** (A) additional examples of in situ staining of old and young GV mice oocytes for constitutive heterochromatin marker H3K9me2. (B) additional examples of in situ staining of old and young GV mice oocytes for facultative heterochromatin marker H3K27me3. (C) additional examples of in situ staining of old and young GV mice oocytes for constitutive heterochromatin marker HP1γ.

**Figure S2: Comparison of heterochromatin staining in SN and NSN configured oocytes:** (A) In situ staining of old and young GV mice oocytes for constitutive heterochromatin marker H3K9me2 in comparison between NSN and SN configuration of nuclei and staining quantification. (B) In situ staining of old and young GV mice oocytes for facultative heterochromatin marker H3K27me3 in comparison between NSN and SN configuration of nuclei and staining quantification. (C) In situ staining of old and young GV mice oocytes for constitutive heterochromatin marker HP1γ in comparison between NSN and SN configuration of nuclei and staining quantification.

**Figure S3: 9 months old oocytes do not lose euchromatin marks (additional examples):** (A) In situ staining of old and young GV mice oocytes, and staining intensity analysis, for the euchromatin marker H3K27Ac (B) Additional examples of in situ staining of old and young GV mice oocytes, and staining intensity analysis, for the euchromatin marker H3K4me3.

**Figure S4: Heterochromatin loss with age in oocytes in a genome-wide phenomenon:** The Ratio between intensely stained loci (A-marked with white circle) and mildly stained loci on the chromosomes was measured (B) and normalized to Hoechst intensity on the same loci. Young and old oocytes show the same ratios. These results show that although these is a significant decrease in heterochromatin signal with age, the ratio between heterochromatin enriched and non-enriched loci on the chromosome is maintained. This means that there is no difference in the amount of heterochromatin decrease along the chromosome.

**Figure S5: Old oocytes show elevated retrotransposon activation and DNA damage (additional examples):** (A) In situ staining of L1- ORF1p in old and young GV mouse oocytes (B) In situ staining of Rad51 in old and young GV mouse oocytes (C) In situ staining of γH2Ax in old and young GV oocytes.

**Figure S6: Analysis of small RNA sequencing in young and old oocytes:** (A) Distribution of RNA fragment sizes in old and young mice. (B) PCA analysis of small RNA sequencing in young and old oocytes. (C) List of the 10 repeat types that are most upregulated with lowest p values in old oocytes (D) List of the 10 repeat types that are most downregulated with lowest p values in old oocytes. In red: LTRs, in green: IAP endogenous retroviruses (ERVs), in yellow: major satellite repeats, in purple: simple repeats

**Figure S7: Old oocytes show elevated retrotransposon RNA processing activity (more examples):** (A) In situ staining for Dicer in old and young GV mouse oocytes. (B) In situ staining for dsRNA in old and young GV mouse oocytes. Note the elevation in signal for dsRNA and Dicer in old oocytes.

**Figure S8: Human oocytes show a reduction of heterochromatin with age:** (A) More examples of in situ staining of GV human oocytes for H3K9me2. (B) More examples of in situ staining of GVBD human oocytes (C) In situ staining of GV human oocytes (N=21) for Rec8. These oocytes show a reduction of signal intensity with age in a linear regression curve (p=0.02). (D) In situ staining of GVBD human oocytes (N=47) for Rec8. These oocytes show a reduction of signal intensity with age in a linear regression curve (p=0.01).

**Figure S9: Treatment of oocytes with Chaetocin mimics natural aging:** (A) chromosome spread MI staining for H3K9me2 in young GV mouse oocytes treated with Chaetocin, showing a reduction in H3K9me2 signal. (B) more examples of in situ staining for dsRNA in young GV mouse oocytes treated with Chaetocin. (C) More examples of in situ staining for L1ORF1p in young GV mouse oocytes treated with Chaetocin.

**Figure S10: Treatment of oocytes with** TSA **mimics natural aging:** (A) Maturation efficiency of young GV mouse oocytes treated with TSA and matured in-vitro (see Methods) (B) In situ staining for H3K27Ac and staining quantification in young GV mouse oocytes treated with TSA showing an elevation in the signal upon treatment. (C) In situ staining for H3K9me2 and staining quantification in young GV mouse oocytes treated with TSA showing a drop in the signal upon treatment. (D) In situ staining for L1ORF1p and staining quantification in young GV mouse oocytes treated with TSA showing an elevation in the nuclear signal upon treatment. (E) In situ staining for dsRNA and staining quantification in young GV mouse oocytes treated with TSA showing an elevation in the signal upon treatment. (F) In situ staining for Rad51 and nuclear localization quantification in young GV mouse oocytes treated with TSA showing an elevation in nuclear localization upon treatment.

**Figure S11: Working model:** Our work has shown that with age, oocytes’ chromatin becomes less and less heterochromatic. The loss of silencing and compaction of heterochromatin results in the loss of transcriptional regulation, enabling the upregulation of retrotransposons. We suggest that the early stages of reproductive aging are the outcome of the gradual loss of repression. The harmful activity of activated retrotransposons eventually leads to the loss of oocytes function, and a reduction in fertility.

## Materials and Methods

### Animals

RCC-C57BL/6JHsd female mice were used for the experiment. For the young group we used 7-10 weeks old mice, and for the old group we used 8-9 months old mice. The experiment was approved by the institutional ethics comity, approval number: MD-19-15938-3. All the mice were housed at the Hebrew University AAALAC-accredited and NIH-accredited SPF facility.

### Mouse Oocyte In-vitro maturation

After Euthanasia, ovaries were collected and dissected in L-15 medium (011151A) supplemented with 200um IBMX (I7018) to prevent meiotic progression. GV oocytes were collected under a binocular using a Stripper (MD-MXL3-STR-CGR). After collection, oocytes were transferred into α-MEM medium (22561021) supplemented with IBMX covered with Mineral oil (M8410-1l) to prevent evaporation, for recovery time of 25 min at 37° in a 5% CO2 incubator, and then washed in IBMX free a MEM medium to initiate meiosis. The oocytes were incubated in α-MEM under oil in the incubator for 6 hours and then examined under a binocular for the presence of a polar body and stained by Hoechst to examine the first meiotic division, or incubated for 18 hours and stained by Hoechst to examine the entry into the second meiotic division metaphase.

### Mouse oocytes collection for in-situ immunofluorescence

After Euthanasia, ovaries were collected and dissected in M2 medium (M7167). GV oocytes were collected under a binocular using a Stripper and washed in hyaluronidase (H4272-30MG) to remove granulosa cells and acidic Tyrode’s Solution (T1788) to remove the Zona Pellucida. The oocytes were fixed using PFA 4% (15710) for 20 min, and then quenched in PBS supplemented with 10mM glycine and 1% BSA.

### Chromosome spreads and immunofluorescence

GV oocytes were collected from ovaries as above and matured to the first meiotic division or second meiosis metaphase as described above. Matured oocytes were washed in M2 medium and then in acidic Tyrode’s Solution to remove the Zona Pellucida. 15 min before the desired time for chromosome spread, oocytes were kept in hypotonic solution, composed of FBS (F7524) diluted in water in 1:1 ratio. Oocytes were then spread in spreading solution (1% PFA buffered to 9.2 pH, supplemented with 0.15% Triton X-100 and 0.03% DDT) on Superfrost plus slides (32090003).

### In situ immunofluorescence

Permeabilization was performed using 0.01% Triton X-100. Cells were cultured in PBS contains 5% BSA for blocking, and first and secondary antibodies were diluted in 0.1 Tween 20 (P9416)/PBS containing 5% BSA. The immune-stained cells were mounted in Vectashield (H-1000) mounting medium containing 80 nM of Hoechst (33342), and sealed on a slide using an Imaging spacer (GBL654002).

### Antibodies

For heterochromatin staining we used antibodies for H3K9me2 (ab1220, concentration of 1:400), H3K27me3 (ab205728, 1:100) HP1γ (ab227478, 1:50). For euchromatin markers we used antibodies for H3K27ac(8173s, 1:100) and H3K4me3(11960s, 1:50). To track retroviral activity and transcription regulation loss we used antibodies for L1-ORF1p (ab216324, 1:100), dsRNA (10020200, 1:500) and Dicer (ab167444, 1:200). Meiotic Cohesin was stained with antibody against REC8 (ab192241 1:200), and DNA damage was assessed using antibody against RAD51(PC130, 1:200), and γH2Ax (0563625UG, 1:500).

### Imaging and quantification

Oocytes were imaged using either Ti-Eclipse Nikon system, with an Andor Zyla nsc05537 camera, or Nikon Yokogawa W1 Spinning Disk, with SCMOS ZYLA camera Every cell was imaged at multiple planes. In order to analyze the images in a quantitative manner, a projection of maximum intensity was created. To measure staining intensity, every assessed region was normalized to an equal-size region in the background (outside the area of interest). The intensity score was generated by dividing the intensity in the area of interest by that of the background. To assess relative localization (for Rad51), the intensities at the different assessed regions were compared and the relative fraction of the signal in the nucleus was calculated. The relative area of the nucleus was also calculated and the fraction of the area was subtracted from the fraction of the signal (% nuclear signal/signal cell- % area-nucleus/area-cell). Hence, the score given is the access localization of Rad51 to the nucleus on top of what was expected by its fraction of the area.

### Small RNA sequencing

#### RNA extraction and amplification

46 oocytes from young and old females (for each replicate) were collected from ovaries as above and dissected in medium M2, and after the granulosa cells and the Zona Pellucida were removed (as mentioned above), the oocytes were inserted into 1 ml of TRIzol (15596026). RNA extraction was done according to the TRIzol reagent user guide, with extraction in 10 μl of nuclease-free water.

Amplification was performed by the SMARTer^®^ smRNA-Seq Kit for Illumina^®^-12 Rxns, TAKARA 635029. Size selection was performed using Agencourt AMPure XP Beads, by adding 50ul of sample to 100ul of beads solution and eluting the DNA bound to the beads. Sequencing was performed at the Ein-Kerem campus interdepartmental equipment park on a NextSeq machine.

#### Trimming and filtering of raw reads

The NextSeq base-calls files were converted to fastq files using the bcl2fastq program with default parameters (without trimming or filtering applied at this stage). Raw reads (fastq files) were inspected for quality issues with FastQC. Following that, and according to the SMARTer smRNA-Seq library construction protocol, the first 3 bases of R1 reads were discarded (positions for template switching), the reads were quality-trimmed at both ends, poly-G sequences (NextSeq’s no signal), adapter sequences, and poly-A sequences were removed from the 3’ end, and finally low quality reads were filtered out. Cutadapt was used for trimming sequences from the end of reads, with parameters that included using a minimal overlap of 1, allowing for read wildcards, and filtering out reads that became shorter than 15 nt. Final filtering by quality was performed using the fastq_quality_filter program of the FASTX package, with a quality threshold of 20 at 90 percent or more of the read’s positions.

#### Alignment and counting

The processed reads were aligned to the mouse transcriptome and genome with TopHat. The genome version was GRCm38, with annotations from Ensembl release 99. The Ensembl genome and gene annotations were supplemented with information for rRNA (adding the 45S pre-rRNA to the genome sequence and annotations), tRNAs (annotations from tRNAscan predictions), and repeat sequences (annotations from the UCSC Genome Browser ‘rmsk’ table). Since some of the external information overlapped existing annotations from the Ensembl original GTF file, those overlapping annotations were discarded. In order to define overlapping annotations, both the start and end positions had to be within 40 bases distance from each other (comparing start position in one source to start position in the other source, and likewise for the end positions). Parameters for the alignment were generally standard except for using -g 1000 in order to allow for all possible alignments of repeat sequences. Quantification was done with htseq-count. Strand information was set to ‘yes’, and each primary alignment was counted. The annotations file had all 5S rRNA genomic locations treated as one gene, had tRNAs counted according to the tRNA type, and repeats counted by their name.

#### Differential expression

Normalization and differential expression analysis were done with the DESeq2 package. Genes with a sum of counts less than 10 over all samples were filtered out, then size factors and dispersion were calculated. Normalized counts were used for several quality control assays, such as counts distributions and principal component analysis, which were calculated and visualized in R. There was a clear batch-effect hence batch-correction was applied. The pair-wise comparison, comparing old to young oocytes, was tested with default parameters, except not using the independent filtering algorithm. In order to find candidate genes for a significant expression difference between young and old oocytes, a baseMean-dependent formula over the log2FoldChange was applied. The formula required a baseMean above 5 and an absolute log2FoldChange bigger than 5/sqrt(baseMean) + 0.6. In addition, the lfcSE had to be low enough, below 1, so signal is repetitive across the biological repeats. Results are detailed in Table S1. We used the calculated p value to filter the best candidate repeat types. Raw data submitted to GEO under accession GSE159789.

#### qRT-PCR

After Euthanasia, ovaries were collected and dissected in M2 medium (M7167) as above. 40 to 60 (number was matched between groups in every experiment) GV oocytes were collected and washed in hyaluronidase (H4272-30MG) to remove granulosa cells, and then washed extensively in M2 until there all the granulosa cells were removed from the oocytes. Oocytes were transferred to a Trizol (15596026) and Chloroform solution for RNA precipitation, treated with RNAse free DNAse (E0013-1D3) to remove DNA and then RNA was purified by using Ampure beads (A63881). Reverse transcription was performed using iScript™ Reverse Transcription Supermix for RT-qPCR (1708891) according to the manufacturer’s recommendations. iTaq™ Universal SYBR^®^ Green Supermix (1725124) was used for amplification reactions. Quantitive PCR measurements were taken using a CFX96 C1000 BioRad machine.

#### Drug treatments

Ovaries were collected and dissected in L-15 medium supplemented with 200um IBMX to prevent meiotic progression as described above. GV oocytes were collected using a Stripper as described above. After collection, oocytes were transferred to α-MEM medium supplemented with IBMX + the desired drug covered with Mineral oil to prevent evaporation, for 4 hours at 37°in a 5% CO2 incubator (Since Chaetocin and IBMX have a cross reactivity with each other, the pause was not performed in the Chaetocin experiments). After the pause oocytes were washed in IBMX-free α-MEM that contains the drug to initiate meiosis. 18.5 hours after oocytes were released from IBMX, oocytes were fixed in PFA, permeabilized, and sealed as described above. Drugs in use were: Trichostatin A (TSA) T1952, Chaetocin (sc-200893).

#### Human oocytes

Human GV oocytes that were retrieved during IVF treatment were incubated for 24h before determination that they remained at this state and did not mature. After informed consent was signed (following IRB approval 0020-16-SZMC) oocytes were treated by piercing of the Zona Pellucida by the embryologists in order to increase permeability. The oocytes were then fixed in 4% PFA the day after retrieval, for 20min at room temperature and then quenched in PBS supplemented with 10mM glycine and 1% BSA. Immunofluorescence was performed as described above.

## References

Bertoldo, M. J., Listijono, D. R., Ho, W. J., Riepsamen, A. H., Goss, D. M., Richani, D., Jin, X. L., Mahbub, S., Campbell, J. M., Habibalahi, A., Loh, W. N., Youngson, N. A., Maniam, J., Wong, A. S. A., Selesniemi, K., Bustamante, S., Li, C., Zhao, Y., Marinova, M. B., Kim, L. J., Lau, L., Wu, R. M., Mikolaizak, A. S., Araki, T., Le Couteur, D. G., Turner, N., Morris, M. J., Walters, K. A., Goldys, E., O’Neill, C., Gilchrist, R. B., Sinclair, D. A., Homer, H. A., & Wu, L. E. (2020, Feb 11). NAD(+) Repletion Rescues Female Fertility during Reproductive Aging. Cell Rep, 30(6), 1670–1681 e1677. https://doi.org/10.1016/j.celrep.2020.01.058

Bonnet-Garnier, A., Feuerstein, P., Chebrout, M., Fleurot, R., Jan, H. U., Debey, P., & Beaujean, N. (2012). Genome organization and epigenetic marks in mouse germinal vesicle oocytes. Int J Dev Biol, 56(10-12), 877–887. https://doi.org/10.1387/ijdb.120149ab

Burkhardt, S., Borsos, M., Szydlowska, A., Godwin, J., Williams, S. A., Cohen, P. E., Hirota, T., Saitou, M., & Tachibana-Konwalski, K. (2016, Mar 7). Chromosome Cohesion Established by Rec8-Cohesin in Fetal Oocytes Is Maintained without Detectable Turnover in Oocytes Arrested for Months in Mice. Curr Biol, 26(5), 678–685. https://doi.org/10.1016/j.cub.2015.12.073

Chen, H., Zheng, X., Xiao, D., & Zheng, Y. (2016, Jun). Age-associated de-repression of retrotransposons in the Drosophila fat body, its potential cause and consequence. Aging Cell, 15(3), 542–552. https://doi.org/10.1111/acel.12465

Chiang, T., Duncan, F. E., Schindler, K., Schultz, R. M., & Lampson, M. A. (2010, Sep 14). Evidence that weakened centromere cohesion is a leading cause of age-related aneuploidy in oocytes. Curr Biol, 20(17), 1522–1528. https://doi.org/10.1016/j.cub.2010.06.069

Crichton, J. H., Dunican, D. S., Maclennan, M., Meehan, R. R., & Adams, I. R. (2014, May). Defending the genome from the enemy within: mechanisms of retrotransposon suppression in the mouse germline. Cell Mol Life Sci, 71(9), 1581–1605. https://doi.org/10.1007/s00018-013-1468-0

De Cecco, M., Criscione, S. W., Peterson, A. L., Neretti, N., Sedivy, J. M., & Kreiling, J. A. (2013, Dec). Transposable elements become active and mobile in the genomes of aging mammalian somatic tissues. Aging (Albany NY), 5(12), 867–883. https://doi.org/10.18632/aging.100621

Dennis, S., Sheth, U., Feldman, J. L., English, K. A., & Priess, J. R. (2012). C. elegans germ cells show temperature and age-dependent expression of Cer1, a Gypsy/Ty3-related retrotransposon. PLoS Pathog, 8(3), e1002591. https://doi.org/10.1371/journal.ppat.1002591

Djeghloul, D., Kuranda, K., Kuzniak, I., Barbieri, D., Naguibneva, I., Choisy, C., Bories, J. C., Dosquet, C., Pla, M., Vanneaux, V., Socie, G., Porteu, F., Garrick, D., & Goodhardt, M. (2016, Jun 14). Age-Associated Decrease of the Histone Methyltransferase SUV39H1 in HSC Perturbs Heterochromatin and B Lymphoid Differentiation. Stem Cell Reports, 6(6), 970–984. https://doi.org/10.1016/j.stemcr.2016.05.007

Dupressoir, A., & Heidmann, T. (1996, Aug). Germ line-specific expression of intracisternal A-particle retrotransposons in transgenic mice. Mol Cell Biol, 16(8), 4495–4503. https://doi.org/10.1128/mcb.16.8.4495

Fernandez-Capetillo, O., Lee, A., Nussenzweig, M., & Nussenzweig, A. (2004, Aug-Sep). H2AX: the histone guardian of the genome. DNA Repair (Amst), 3(8-9), 959–967. https://doi.org/10.1016/j.dnarep.2004.03.024

Flemr, M., Malik, R., Franke, V., Nejepinska, J., Sedlacek, R., Vlahovicek, K., & Svoboda, P. (2013, Nov 7). A retrotransposon-driven dicer isoform directs endogenous small interfering RNA production in mouse oocytes. Cell, 155(4), 807–816. https://doi.org/10.1016/j.cell.2013.10.001

Gasior, S. L., Wakeman, T. P., Xu, B., & Deininger, P. L. (2006, Apr 14). The human LINE-1 retrotransposon creates DNA double-strand breaks. J Mol Biol, 357(5), 1383–1393. https://doi.org/10.1016/j.jmb.2006.01.089

Gildemeister, O. S., Sage, J. M., & Knight, K. L. (2009, Nov 13). Cellular redistribution of Rad51 in response to DNA damage: novel role for Rad51C. J Biol Chem, 284(46), 31945–31952. https://doi.org/10.1074/jbc.M109.024646

Greiner, D., Bonaldi, T., Eskeland, R., Roemer, E., & Imhof, A. (2005, Aug). Identification of a specific inhibitor of the histone methyltransferase SU(VAR)3-9. Nat Chem Biol, 1(3), 143–145. https://doi.org/10.1038/nchembio721

Gruhn, J. R., Zielinska, A. P., Shukla, V., Blanshard, R., Capalbo, A., Cimadomo, D., Nikiforov, D., Chan, A. C., Newnham, L. J., Vogel, I., Scarica, C., Krapchev, M., Taylor, D., Kristensen, S. G., Cheng, J., Ernst, E., Bjorn, A. B., Colmorn, L. B., Blayney, M., Elder, K., Liss, J., Hartshorne, G., Grondahl, M. L., Rienzi, L., Ubaldi, F., McCoy, R., Lukaszuk, K., Andersen, C. Y., Schuh, M., & Hoffmann, E. R. (2019, Sep 27). Chromosome errors in human eggs shape natural fertility over reproductive life span. Science, 365(6460), 1466–1469. https://doi.org/10.1126/science.aav7321

Hanna, C. W., Demond, H., & Kelsey, G. (2018, Sep 1). Epigenetic regulation in development: is the mouse a good model for the human? Hum Reprod Update, 24(5), 556–576. https://doi.org/10.1093/humupd/dmy021

Hassold, T., & Hunt, P. (2001, Apr). To err (meiotically) is human: the genesis of human aneuploidy. Nat Rev Genet, 2(4), 280–291. https://doi.org/10.1038/35066065

Igarashi, H., Takahashi, T., & Nagase, S. (2015, Oct). Oocyte aging underlies female reproductive aging: biological mechanisms and therapeutic strategies. Reprod Med Biol, 14(4), 159–169. https://doi.org/10.1007/s12522-015-0209-5

Jeon, H. J., Kim, Y. S., Kim, J. G., Heo, K., Pyo, J. H., Yamaguchi, M., Park, J. S., & Yoo, M. A. (2018, Jul). Effect of heterochromatin stability on intestinal stem cell aging in Drosophila. Mech Ageing Dev, 173, 50–60. https://doi.org/10.1016/j.mad.2018.04.001

Jin, Y. X., Zhao, M. H., Zheng, Z., Kwon, J. S., Lee, S. K., Cui, X. S., & Kim, N. H. (2014). Histone deacetylase inhibitor trichostatin A affects porcine oocyte maturation in vitro. Reprod Fertil Dev, 26(6), 806–816. https://doi.org/10.1071/RD13013

Kageyama, S., Liu, H., Kaneko, N., Ooga, M., Nagata, M., & Aoki, F. (2007, Jan). Alterations in epigenetic modifications during oocyte growth in mice. Reproduction, 133(1), 85–94. https://doi.org/10.1530/REP-06-0025

Keenan, C. R., Iannarella, N., Naselli, G., Bediaga, N. G., Johanson, T. M., Harrison, L. C., & Allan, R. S. (2020, Jun 4). Extreme disruption of heterochromatin is required for accelerated hematopoietic aging. Blood, 135(23), 2049–2058. https://doi.org/10.1182/blood.2019002990

Koehler, K. E., Schrump, S. E., Cherry, J. P., Hassold, T. J., & Hunt, P. A. (2006, Aug 8). Near-human aneuploidy levels in female mice with homeologous chromosomes. Curr Biol, 16(15), R579–580. https://doi.org/10.1016/j.cub.2006.07.018

Lister, L. M., Kouznetsova, A., Hyslop, L. A., Kalleas, D., Pace, S. L., Barel, J. C., Nathan, A., Floros, V., Adelfalk, C., Watanabe, Y., Jessberger, R., Kirkwood, T. B., Hoog, C., & Herbert, M. (2010, Sep 14). Age-related meiotic segregation errors in mammalian oocytes are preceded by depletion of cohesin and Sgo2. Curr Biol, 20(17), 1511–1521. https://doi.org/10.1016/j.cub.2010.08.023

Liu, L., & Keefe, D. L. (2008, Jan). Defective cohesin is associated with age-dependent misaligned chromosomes in oocytes. Reprod Biomed Online, 16(1), 103–112. https://doi.org/10.1016/s1472-6483(10)60562-7

Lopez-Otin, C., Blasco, M. A., Partridge, L., Serrano, M., & Kroemer, G. (2013, Jun 6). The hallmarks of aging. Cell, 153(6), 1194–1217. https://doi.org/10.1016/j.cell.2013.05.039

Malki, S., van der Heijden, G. W., O’Donnell, K. A., Martin, S. L., & Bortvin, A. (2014, Jun 9). A role for retrotransposon LINE-1 in fetal oocyte attrition in mice. Dev Cell, 29(5), 521–533. https://doi.org/10.1016/j.devcel.2014.04.027

Manosalva, I., & Gonzalez, A. (2010, Dec). Aging changes the chromatin configuration and histone methylation of mouse oocytes at germinal vesicle stage. Theriogenology, 74(9), 1539–1547. https://doi.org/10.1016/j.theriogenology.2010.06.024

Merriman, J. A., Jennings, P. C., McLaughlin, E. A., & Jones, K. T. (2012, Feb). Effect of aging on superovulation efficiency, aneuploidy rates, and sister chromatid cohesion in mice aged up to 15 months. Biol Reprod, 86(2), 49. https://doi.org/10.1095/biolreprod.111.095711

Miao, Y. L., Kikuchi, K., Sun, Q. Y., & Schatten, H. (2009, Sep-Oct). Oocyte aging: cellular and molecular changes, developmental potential and reversal possibility. Hum Reprod Update, 15(5), 573–585. https://doi.org/10.1093/humupd/dmp014

Nagaoka, S. I., Hassold, T. J., & Hunt, P. A. (2012, Jun 18). Human aneuploidy: mechanisms and new insights into an age-old problem. Nat Rev Genet, 13(7), 493–504. https://doi.org/10.1038/nrg3245

Naruse, C., Abe, K., Yoshihara, T., Kato, T., Nishiuchi, T., & Asano, M. (2020, Mar). Heterochromatin protein 1gamma deficiency decreases histone H3K27 methylation in mouse neurosphere neuronal genes. FASEB J, 34(3), 3956–3968. https://doi.org/10.1096/fj.201900139R

Patterson, M. N., Scannapieco, A. E., Au, P. H., Dorsey, S., Royer, C. A., & Maxwell, P. H. (2015, Oct). Preferential retrotransposition in aging yeast mother cells is correlated with increased genome instability. DNA Repair (Amst), 34, 18–27. https://doi.org/10.1016/j.dnarep.2015.07.004

Soifer, H. S., Zaragoza, A., Peyvan, M., Behlke, M. A., & Rossi, J. J. (2005). A potential role for RNA interference in controlling the activity of the human LINE-1 retrotransposon. Nucleic Acids Res, 33(3), 846–856. https://doi.org/10.1093/nar/gki223

Svobodova, E., Kubikova, J., & Svoboda, P. (2016, Jun). Production of small RNAs by mammalian Dicer. Pflugers Arch, 468(6), 1089–1102. https://doi.org/10.1007/s00424-016-1817-6

Tarallo, V., Hirano, Y., Gelfand, B. D., Dridi, S., Kerur, N., Kim, Y., Cho, W. G., Kaneko, H., Fowler, B. J., Bogdanovich, S., Albuquerque, R. J., Hauswirth, W. W., Chiodo, V. A., Kugel, J. F., Goodrich, J. A., Ponicsan, S. L., Chaudhuri, G., Murphy, M. P., Dunaief, J. L., Ambati, B. K., Ogura, Y., Yoo, J. W., Lee, D. K., Provost, P., Hinton, D. R., Nunez, G., Baffi, J. Z., Kleinman, M. E., & Ambati, J. (2012, May 11). DICER1 loss and Alu RNA induce age-related macular degeneration via the NLRP3 inflammasome and MyD88. Cell, 149(4), 847–859. https://doi.org/10.1016/j.cell.2012.03.036

Tharp, M. E., Malki, S., & Bortvin, A. (2020, Jan 16). Maximizing the ovarian reserve in mice by evading LINE-1 genotoxicity. Nat Commun, 11(1), 330. https://doi.org/10.1038/s41467-019-14055-8

Trelogan, S. A., & Martin, S. L. (1995, Feb 28). Tightly regulated, developmentally specific expression of the first open reading frame from LINE-1 during mouse embryogenesis. Proc Natl Acad Sci U S A, 92(5), 1520–1524. https://doi.org/10.1073/pnas.92.5.1520

Tsutsumi, M., Fujiwara, R., Nishizawa, H., Ito, M., Kogo, H., Inagaki, H., Ohye, T., Kato, T., Fujii, T., & Kurahashi, H. (2014). Age-related decrease of meiotic cohesins in human oocytes. PLoS One, 9(5), e96710. https://doi.org/10.1371/journal.pone.0096710

Wang, H. Y., Long, Q. Y., Tang, S. B., Xiao, Q., Gao, C., Zhao, Q. Y., Li, Q. L., Ye, M., Zhang, L., Li, L. Y., & Wu, M. (2019, Mar 18). Histone demethylase KDM3A is required for enhancer activation of hippo target genes in colorectal cancer. Nucleic Acids Res, 47(5), 2349–2364. https://doi.org/10.1093/nar/gky1317

Yoshida, M., Matsuyama, A., Komatsu, Y., & Nishino, N. (2003, Nov). From discovery to the coming generation of histone deacetylase inhibitors. Curr Med Chem, 10(22), 2351–2358. https://doi.org/10.2174/0929867033456602

Yue, M. X., Fu, X. W., Zhou, G. B., Hou, Y. P., Du, M., Wang, L., & Zhu, S. E. (2012, Jul). Abnormal DNA methylation in oocytes could be associated with a decrease in reproductive potential in old mice. J Assist Reprod Genet, 29(7), 643–650. https://doi.org/10.1007/s10815-012-9780-4

Zhang, W., Li, J., Suzuki, K., Qu, J., Wang, P., Zhou, J., Liu, X., Ren, R., Xu, X., Ocampo, A., Yuan, T., Yang, J., Li, Y., Shi, L., Guan, D., Pan, H., Duan, S., Ding, Z., Li, M., Yi, F., Bai, R., Wang, Y., Chen, C., Yang, F., Li, X., Wang, Z., Aizawa, E., Goebl, A., Soligalla, R. D., Reddy, P., Esteban, C. R., Tang, F., Liu, G. H., & Belmonte, J. C. (2015, Jun 5). Aging stem cells. A Werner syndrome stem cell model unveils heterochromatin alterations as a driver of human aging. Science, 348(6239), 1160–1163. https://doi.org/10.1126/science.aaa1356

