## Supplementary material for "Loss of heterochromatin and retrotransposon silencing constitute an early phase in oocyte aging": S11

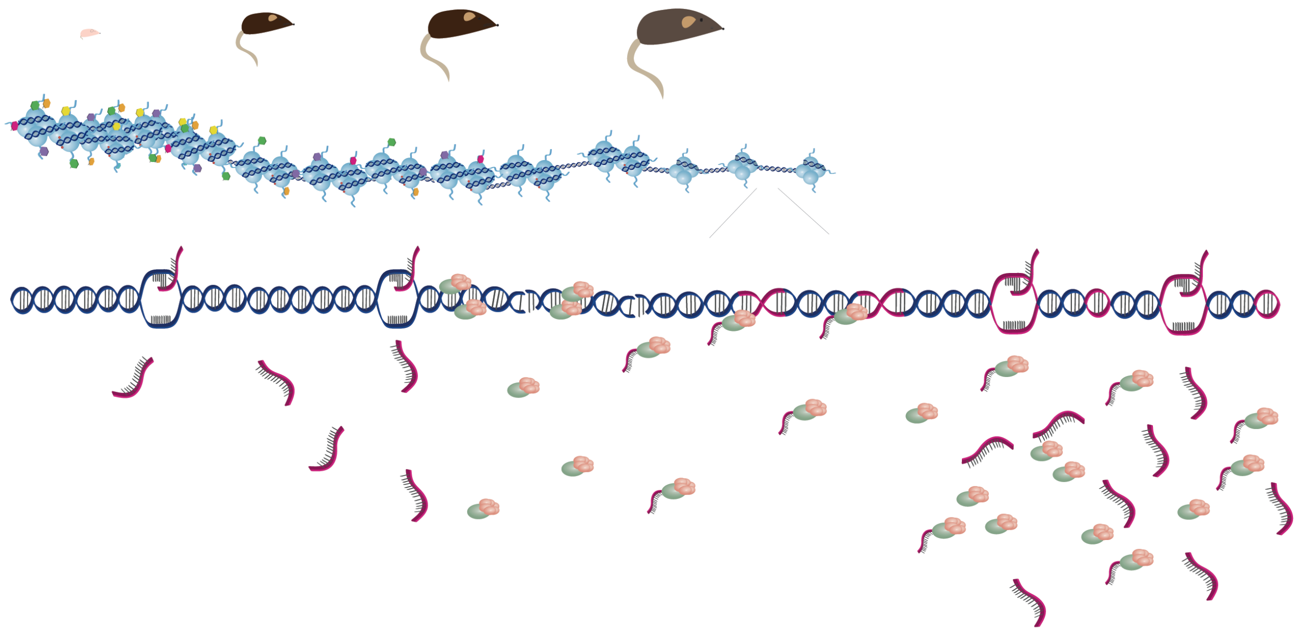

Figure S11: Working model: Our work has shown that with age, oocytes' chromatin becomes less and less heterochromatic. The loss of silencing and compaction of heterochromatin results in the loss of transcriptional regulation , enabling the upregulation of retrotransposons. We suggest that the early stages of reproductive aging are the outcome of the gradual loss of repression. The harmful activity
