## Supplementary material for "Loss of heterochromatin and retrotransposon silencing constitute an early phase in oocyte aging": S10

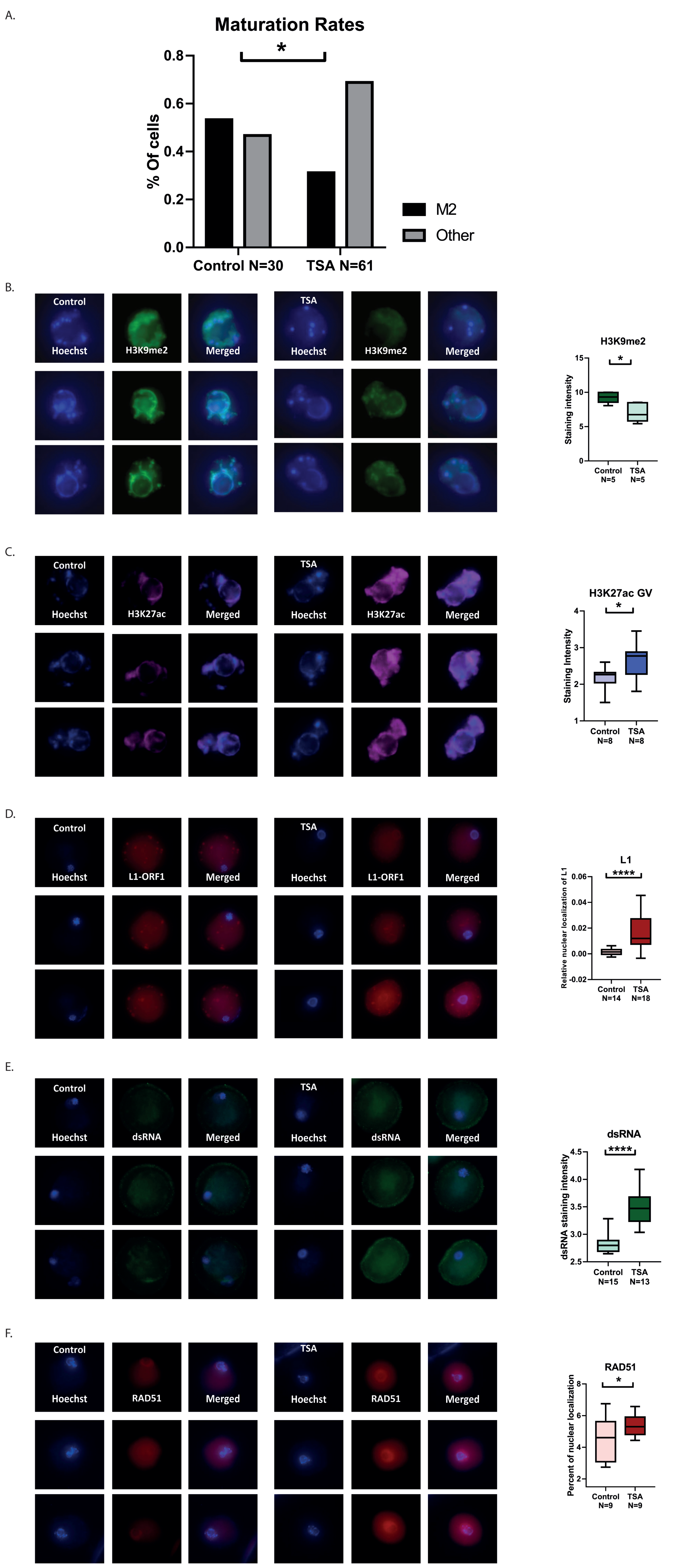


Figure S10: Treatment of oocytes with TSA mimics natural aging: (A) Maturation efficiency of young GV mouse oocytes treated with TSA and matured in-vitro (see Methods) (B) In situ staining for H3K27Ac and staining quantification in young GV mouse oocytes treated with TSA showing an elevation in the signal upon treatment. (C) In situ staining for H3K9me2 and staining quantification in young GV mouse oocytes treated with TSA showing a drop in the signal upon treatment. (D) In situ staining for L1ORF1p and staining quantification in young GV mouse oocytes treated with TSA showing an elevation in the nuclear signal upon treatment. (E) In situ staining for dsRNA and staining quantification in young GV mouse oocytes treated with TSA showing an elevation in the signal upon treatment. (F) In situ staining for Rad51 and nuclear localization quantification in young GV mouse oocytes treated with TSA showing an elevation in nuclear localization upon treatment.
