## Supplementary material for "Loss of heterochromatin and retrotransposon silencing constitute an early phase in oocyte aging": S9

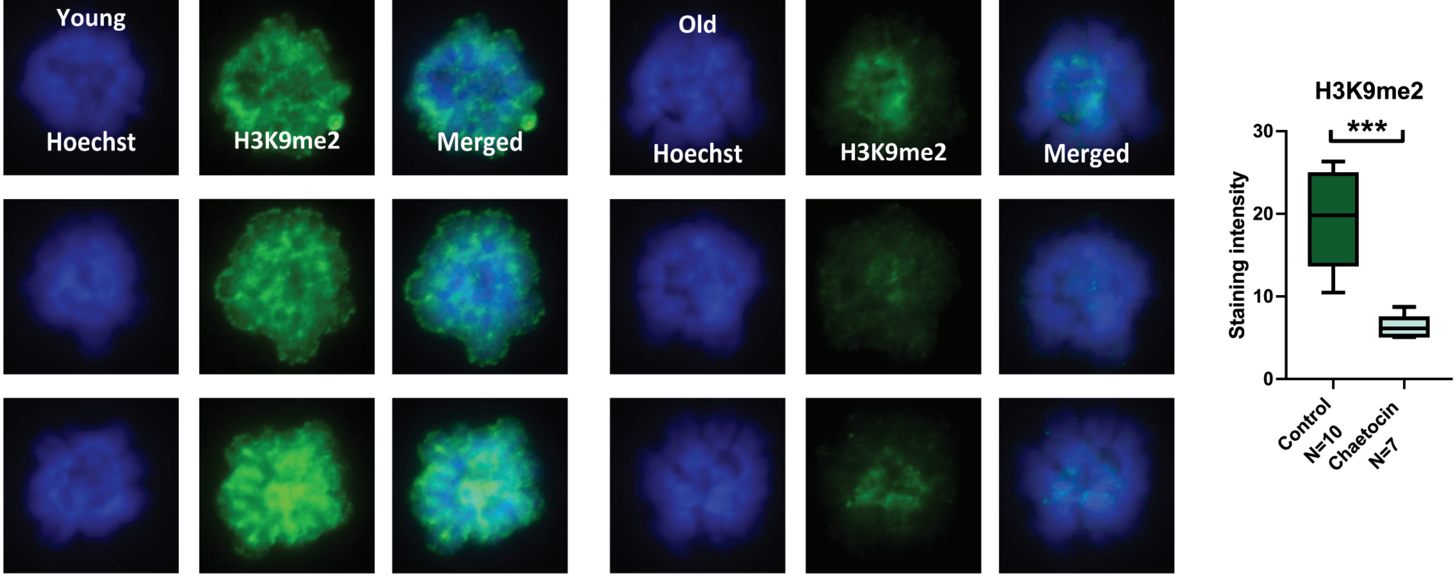


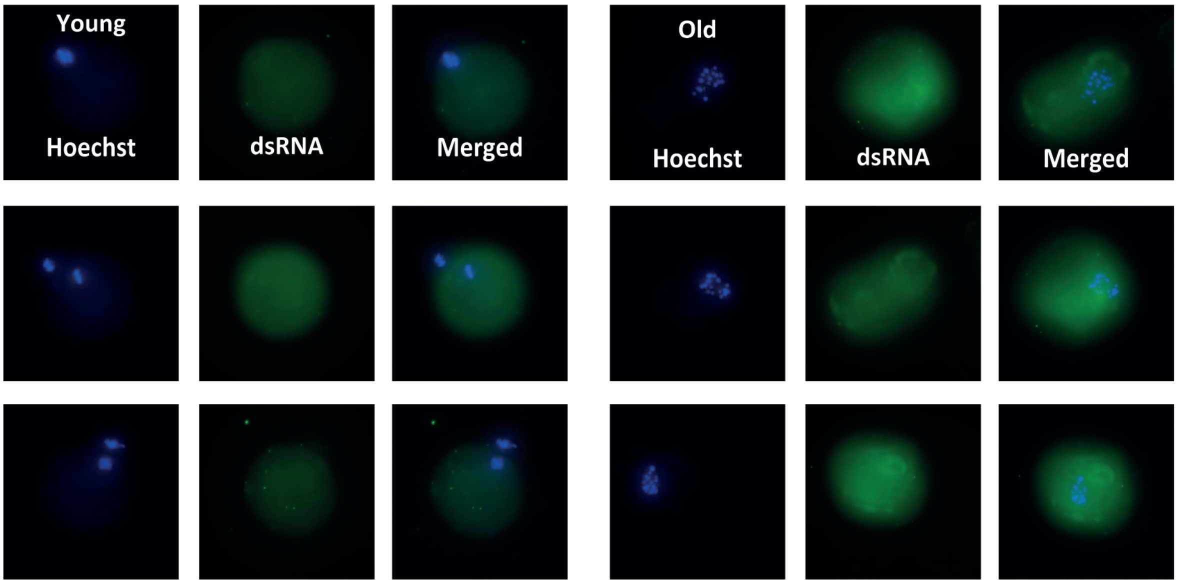


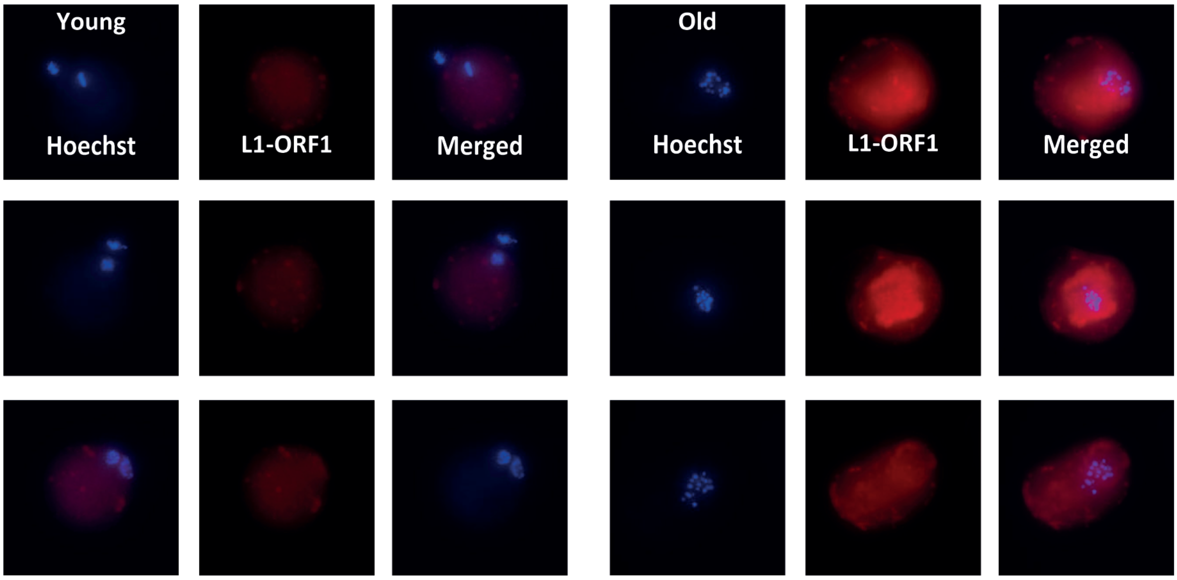


Figure S9: Treatment of oocytes with Chaetocin mimics natural aging: (A) chromosome spread MI staining for H3K9me2 in young GV mouse oocytes treated with Chaetocin, showing a reduction in H3K9me2 signal . (B) more examples of in situ staining for dsRNA in young GV mouse oocytes treated with Chaetocin. (C) More examples of in situ staining for L1ORF1p in young GV mouse oocytes treated with Chaetocin.
