## Supplementary material for "Loss of heterochromatin and retrotransposon silencing constitute an early phase in oocyte aging": S8

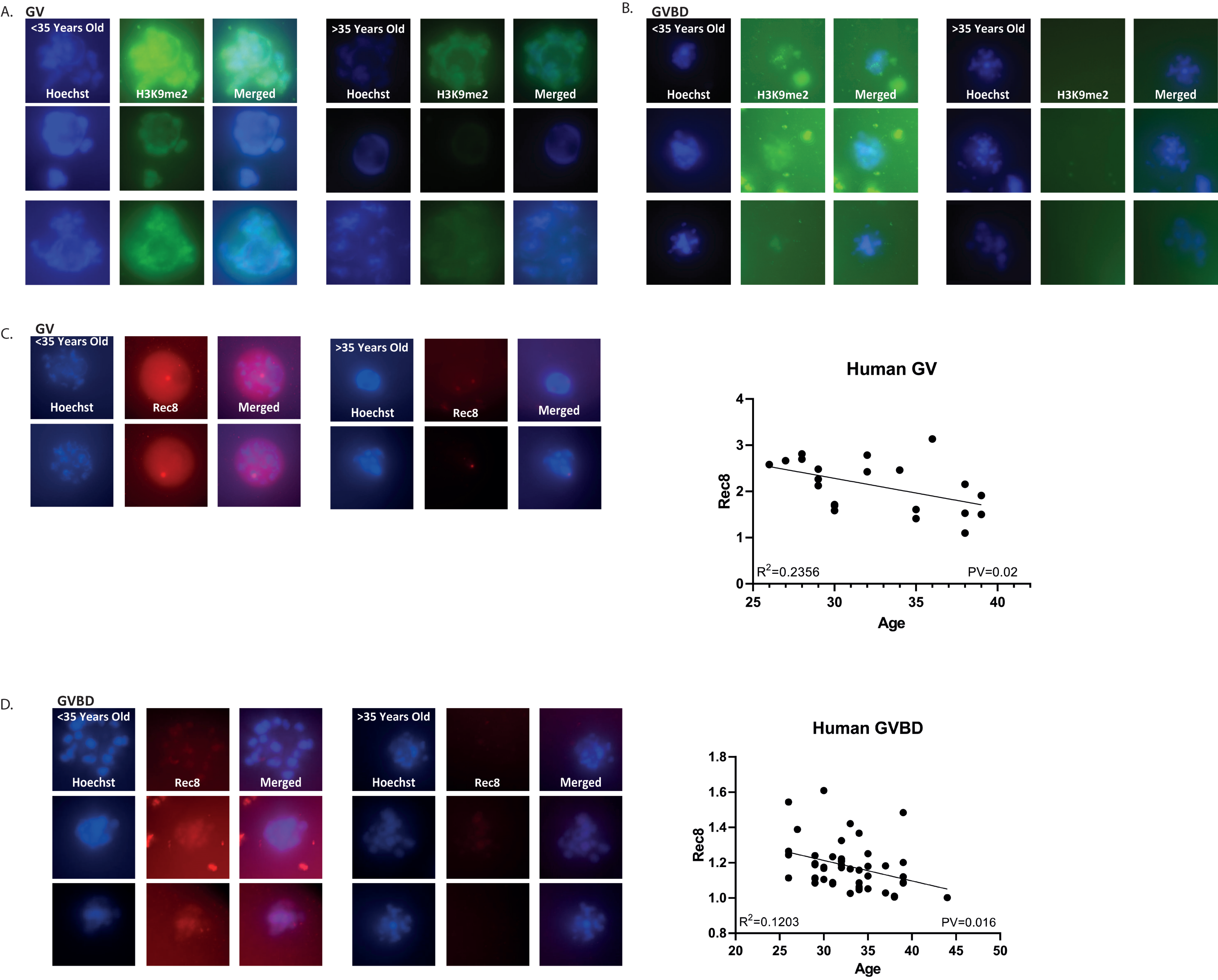


Figure S8: Human oocytes show a reduction of heterochromatin with age: (A) More examples of in situ staining of GV human oocytes for H3K9me2. (B) More examples of in situ staining of GVBD human oocytes (C) In situ staining of GV human oocytes (N =21) for Rec8. These oocytes show a reduction of signal intensity with age in a linear regression curve (p=0.02). (D) In situ staining of GVBD human oocytes (N =47) for Rec8. These oocytes show a reduction of signal intensity with age in a linear regression curve (p=0.01).
