## Supplementary material for "Loss of heterochromatin and retrotransposon silencing constitute an early phase in oocyte aging": S7

A.

B.

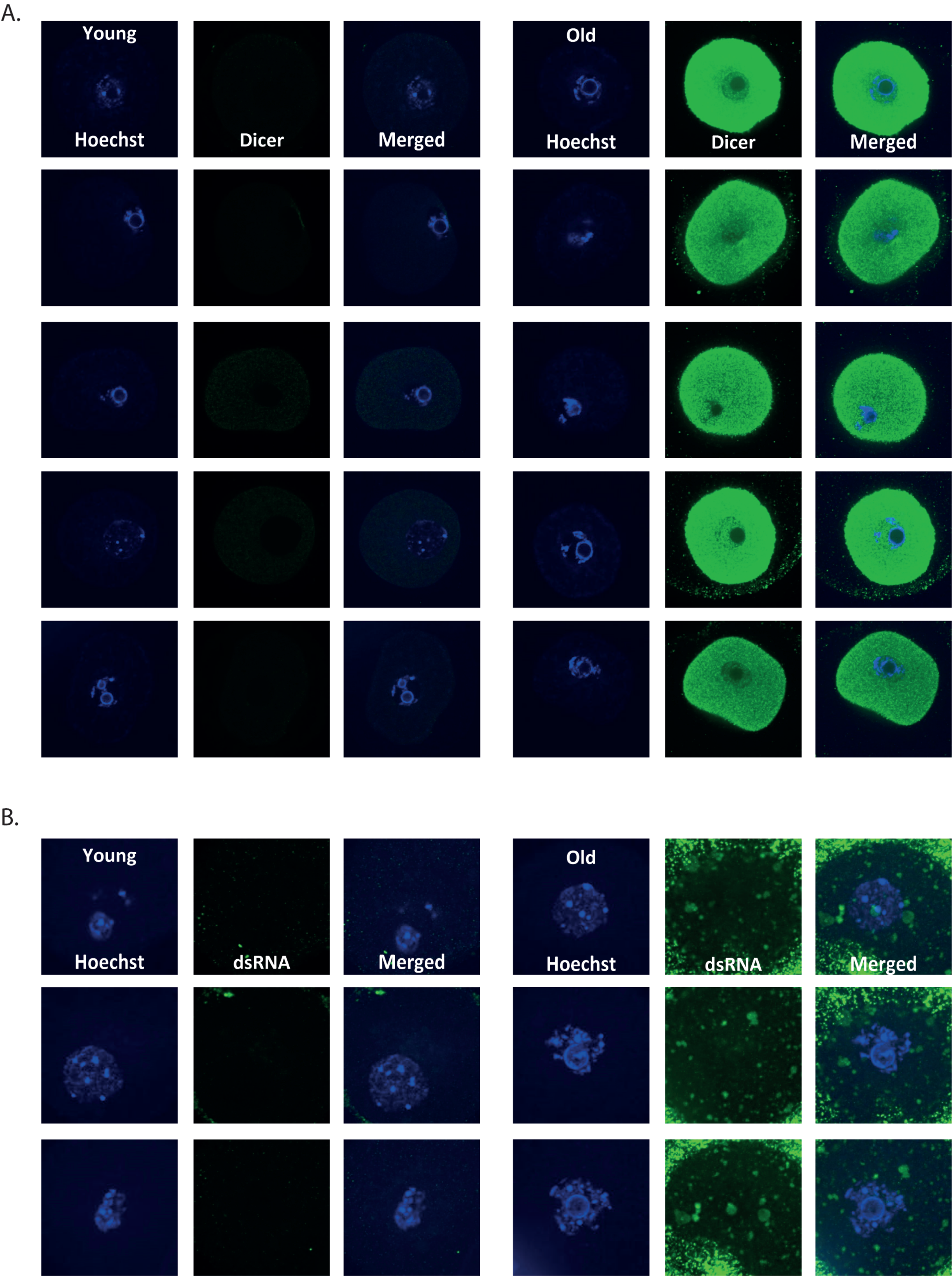


Figure S7: Old oocytes show elevated retrotransposon RNA processing activity (more examples): (A) In situ staining for Dicer in old and young GV mouse oocytes. (B) In situ staining for dsRNA in old and young GV mouse oocytes. Note the elevation in signal for dsRNA and Dicer in old oocytes.
