## Supplementary material for "Loss of heterochromatin and retrotransposon silencing constitute an early phase in oocyte aging": S6

A.

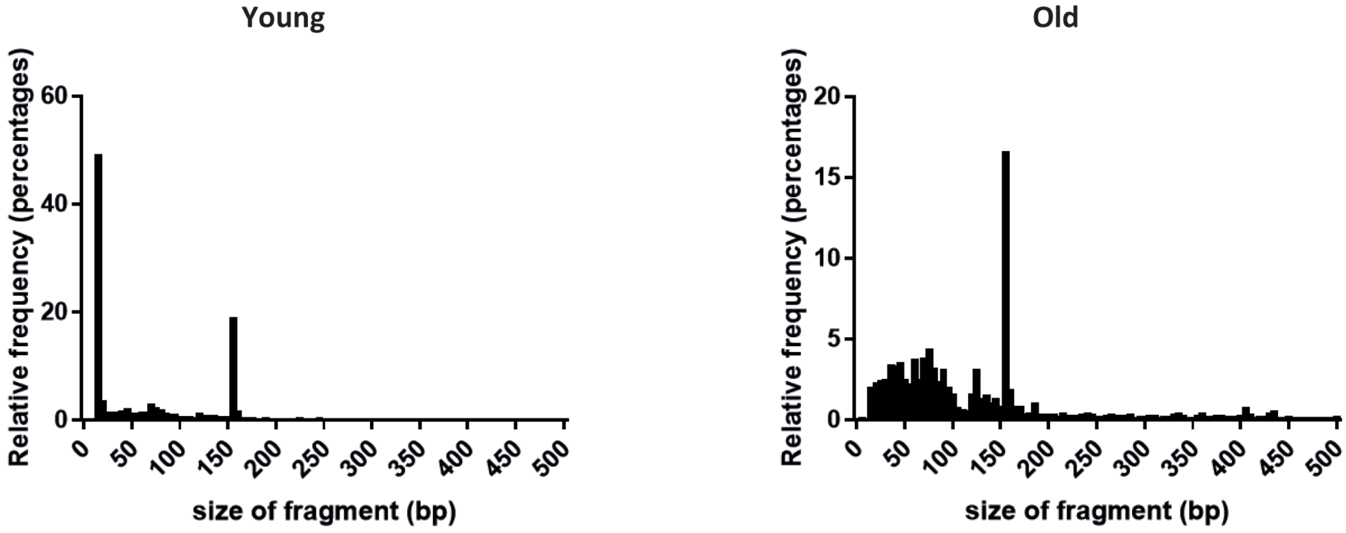


B.

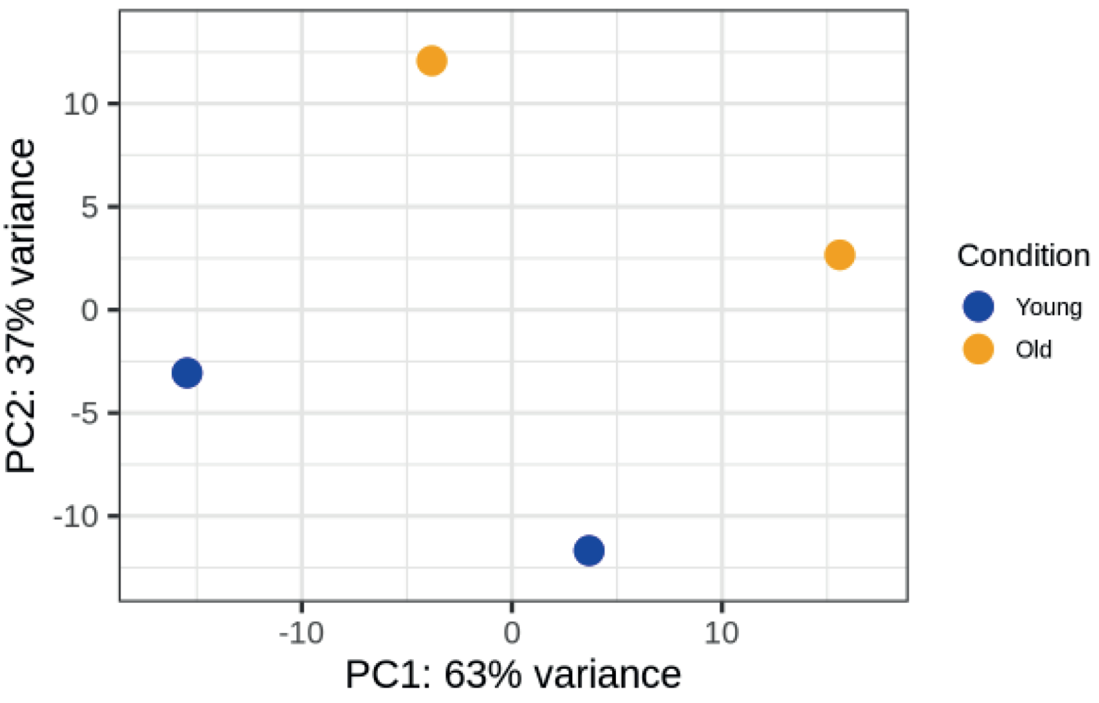


C.

| Gene name | Log2 Fold change O/Y | P value |
|---|---|---|
| LTRIS_Mm | 1.636192813 | 0.003857471 |
| LTR16B | 3.146059483 | 0.014051912 |
| RMER10A | 1.304612379 | 0.017098639 |
| LTRIS_Mus | 1.237043185 | 0.04048463 |
| IAPEz-int | 0.808511756 | 0.073505118 |
| IAPLTR1a_Mm | 0.910646363 | 0.076982915 |
| ORR1F | 0.86968009 | 0.103785775 |
| MT2_Mm | 0.729528591 | 0.144445027 |
| RLTR10F | 1.834640022 | 0.164259341 |
| LTR67B | 1.210938376 | 0.168926367 |

D.

| Gene name | Log2 Fold change O/Y | P value |
|---|---|---|
| RLTR20A4 | -1.708851982 | 0.000581416 |
| GSAT_MM | -1.925634962 | 0.00732218 |
| RLTR24 | -1.645299015 | 0.009572431 |
| MLTR18_MM | -1.751266905 | 0.016515769 |
| (CCAA)n | -2.433041108 | 0.01787433 |
| (TAGA)n | -3.724388175 | 0.029904941 |
| L1MDa | -1.044055494 | 0.051350652 |
| FordPrefect | -1.255606461 | 0.065418401 |
| RMER4B | -0.969625575 | 0.071672173 |
| L1MB3 | -0.829939655 | 0.079516611 |

E.

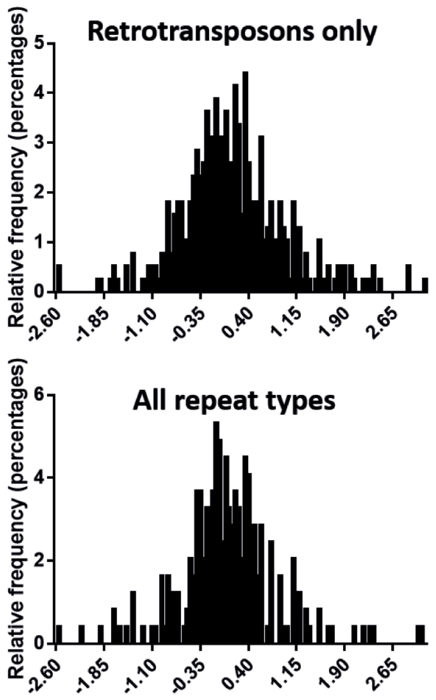


Figure S6: Analysis of small RNA sequencing in young and old oocytes: (A) Distribution of RNA fragment sizes in old and young mice. (B) PCA analysis of small RNA sequencing in young and old oocytes. (C) List of the 10 repeat types that are most upregulated with lowest p values in old oocytes (D) List of the 10 repeat types that are most downregulated with lowest p values in old oocytes. In red: LTRs, in green: IAP endogenous retroviruses (ERVs), in yellow: major satellite repeats, in purple: simple repeats. (E) Deviesion of log-fold change in expression in old oocytes compareto young, of retrotransposon (top) the compared to all the repeats (bottom)
