## Supplementary material for "Loss of heterochromatin and retrotransposon silencing constitute an early phase in oocyte aging": S5

A.

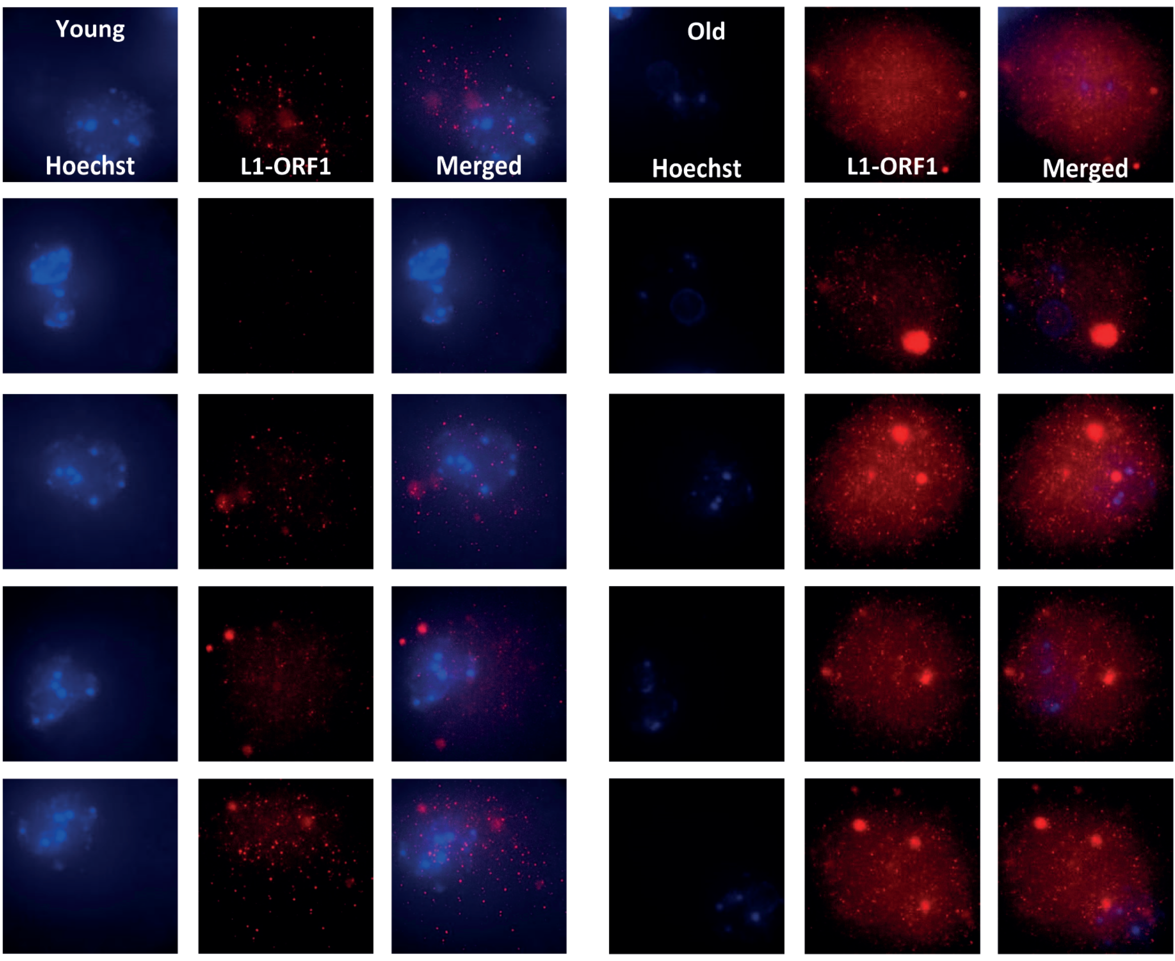


B.

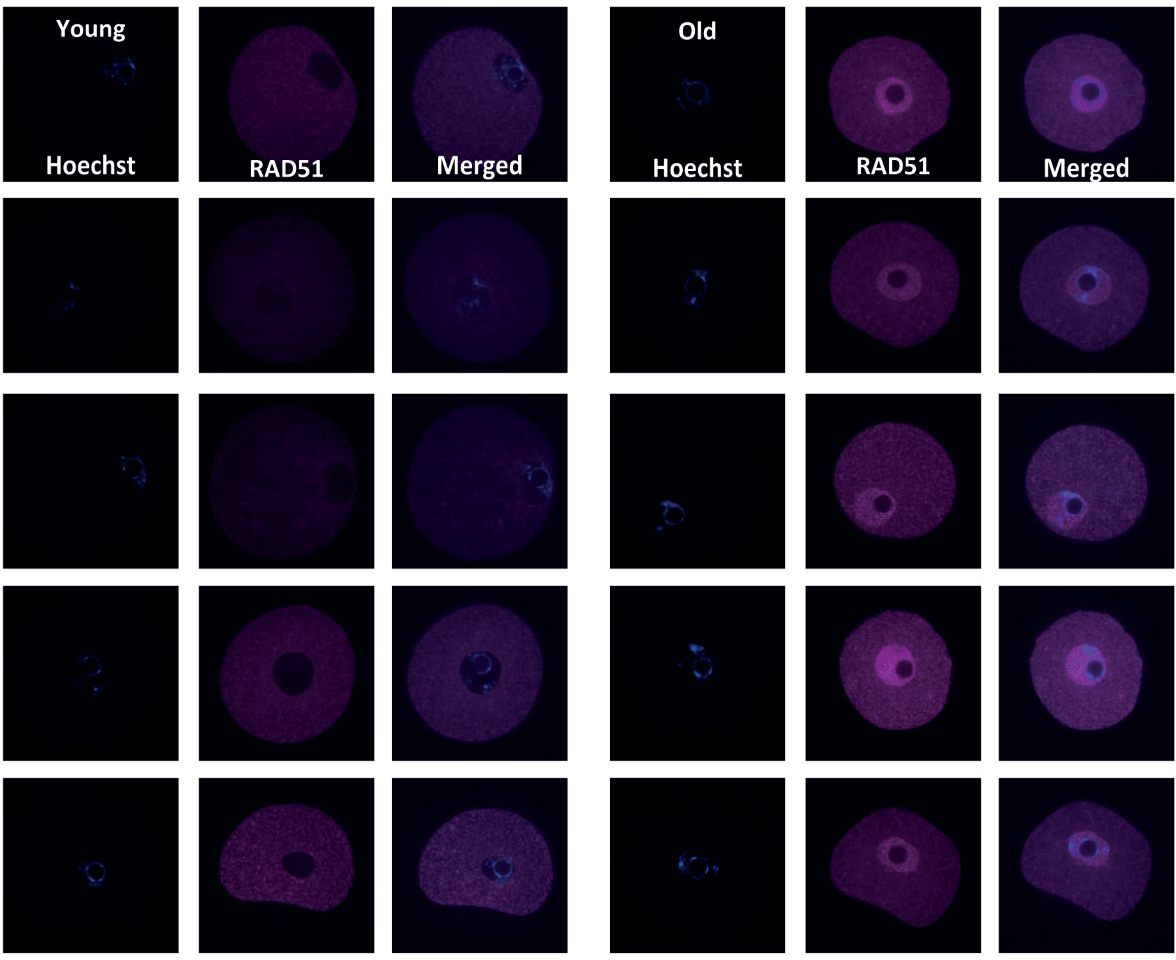


C.

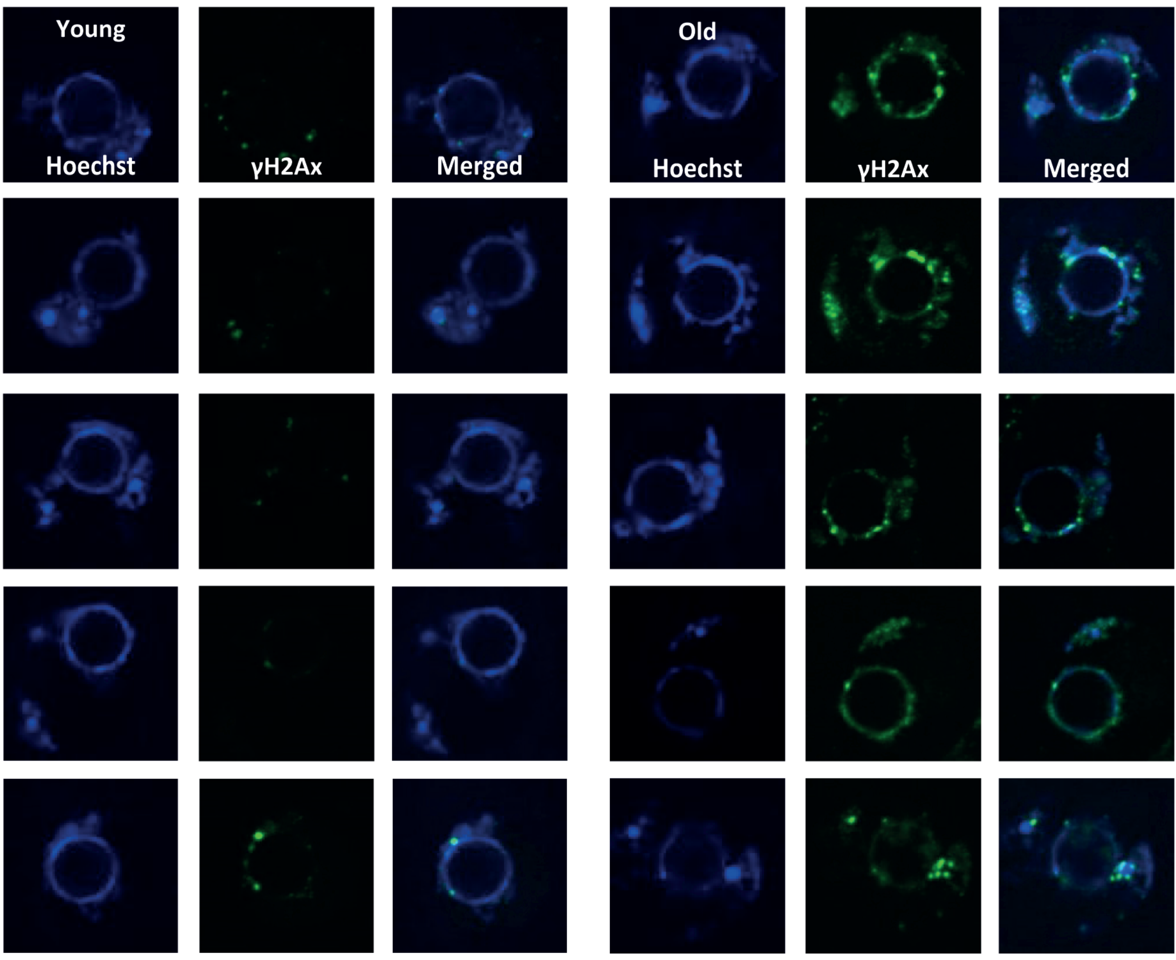


Figure S5: Old oocytes show elevated retrotransposon activation and DNA damage (additional examples): (A) In situ staining of L1- ORF1p in old and young GV mouse oocytes (B) In situ staining of Rad51 in old and young GV mouse oocytes (c) In situ staining of γH2Ax in old and young GV oocytes.
