## Supplementary material for "Loss of heterochromatin and retrotransposon silencing constitute an early phase in oocyte aging": S4

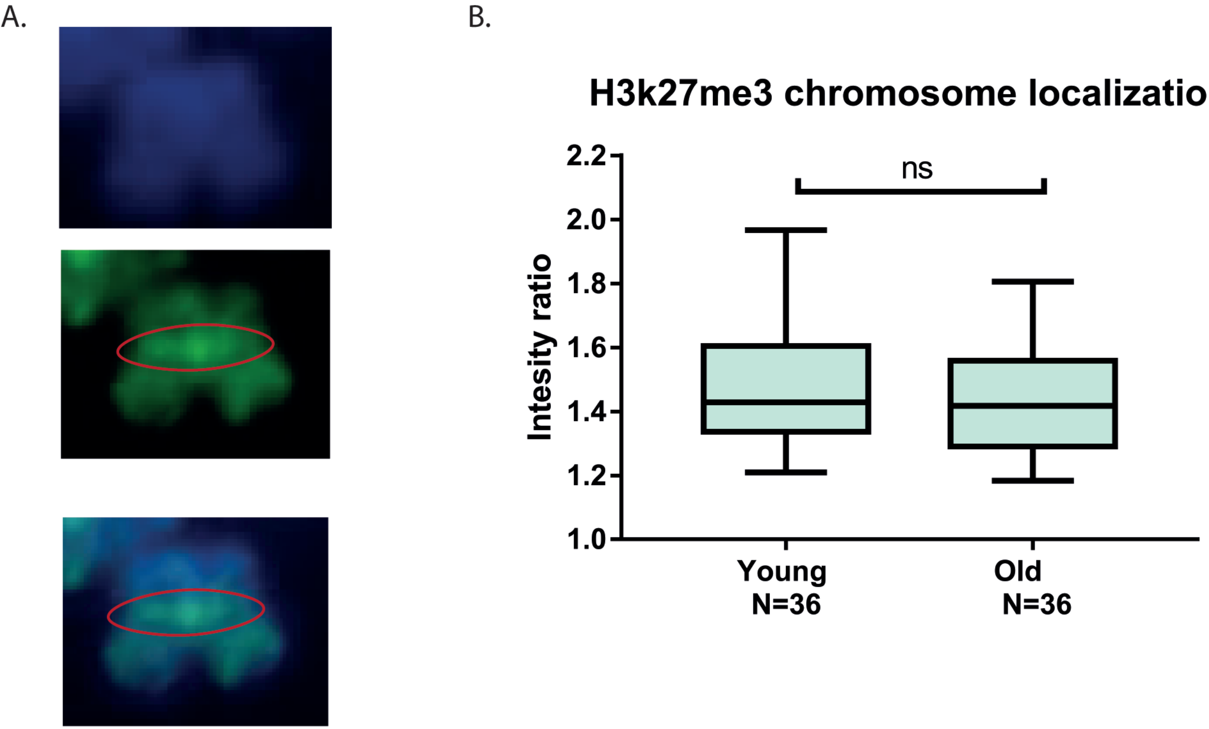


Figure S4: Heterochromatin loss with age in oocytes in a genome-wide phenomenon: The Ratio between intensely stained loci (A-marked with white circle) and mildly stained loci on the chromosomes was measured (B) and normalized to Hoechst intensity on the same loci. Young and old oocytes show the same ratios. These results show that although these is a significant decrease in heterochromatin signal with age, the ratio between heterochromatin enriched and non-enriched loci on the chromosome is maintained. This means that there is no difference in the amount of heterochromatin decrease along the chromosome.
