## Supplementary material for "Loss of heterochromatin and retrotransposon silencing constitute an early phase in oocyte aging": S3

A.

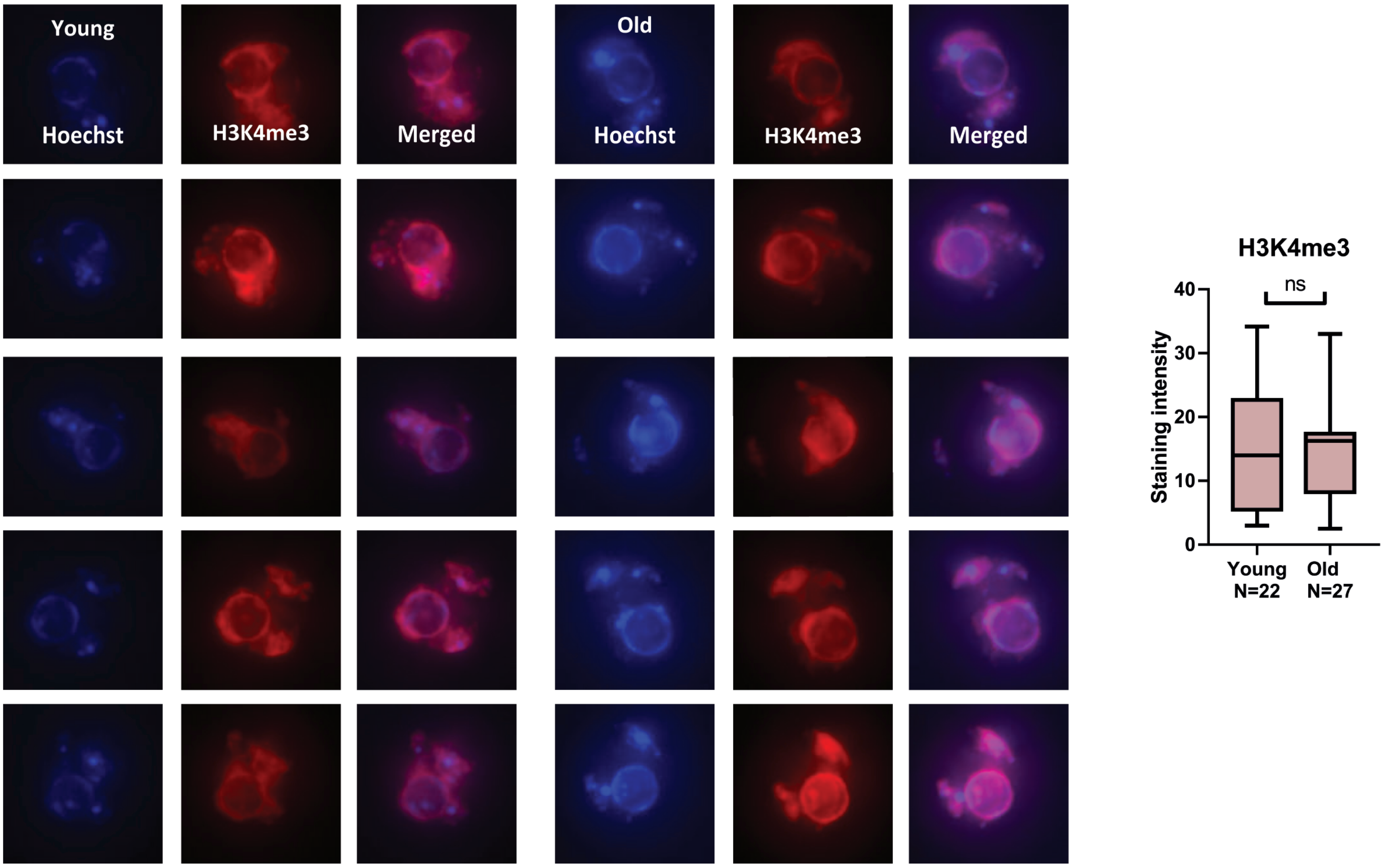


B.

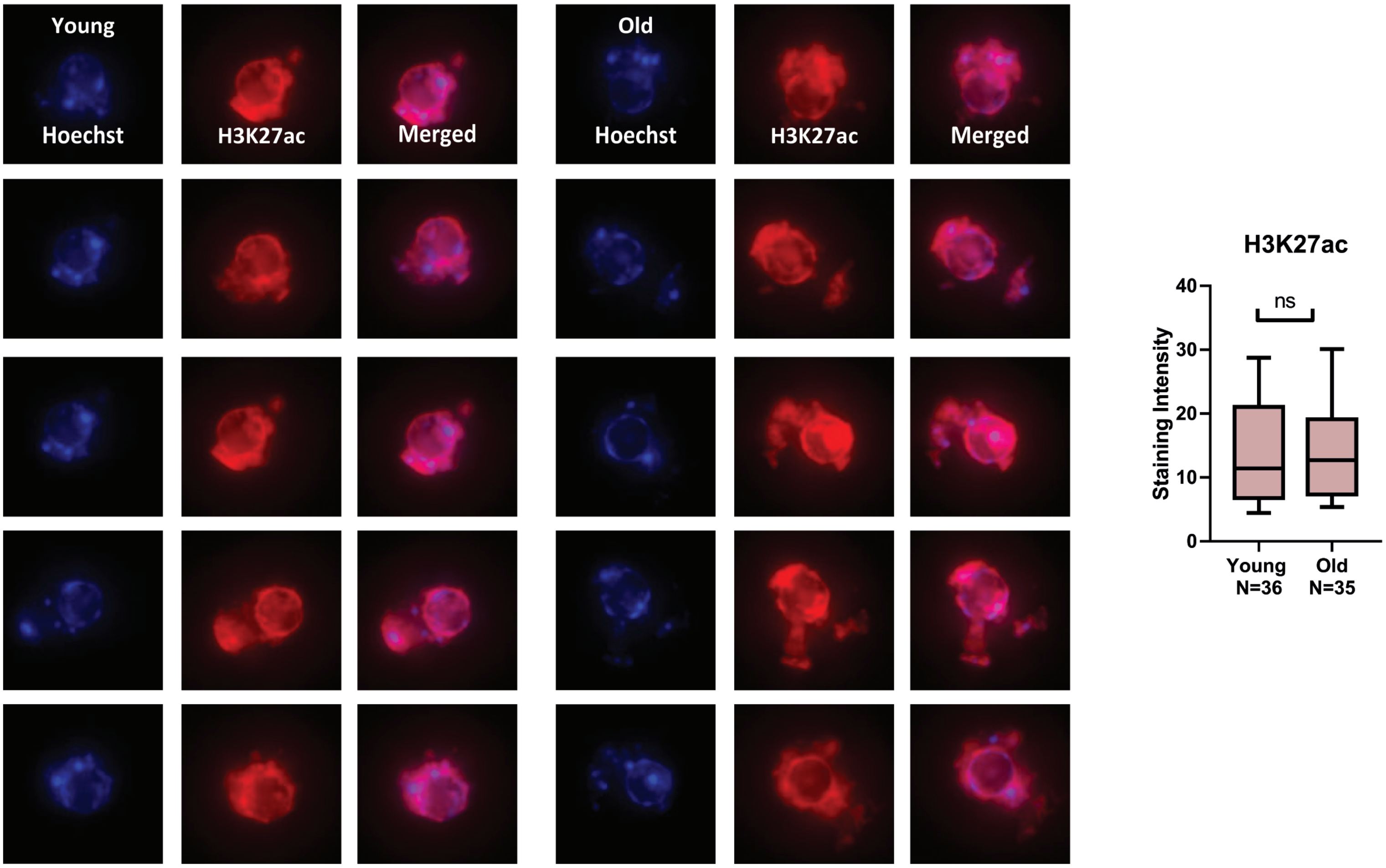


Figure S3: 9 months old oocytes do not lose euchromatin marks (additional examples): (A) In situ staining of old and young GV mice oocytes, and staining intensity analysis, for the euchromatin marker H3K4me3 (B) Additional examples of in situ staining of old and young GV mice oocytes, and staining intensity analysis, for the euchromatin marker H3K27Ac.
