## Supplementary material for "Loss of heterochromatin and retrotransposon silencing constitute an early phase in oocyte aging": S2

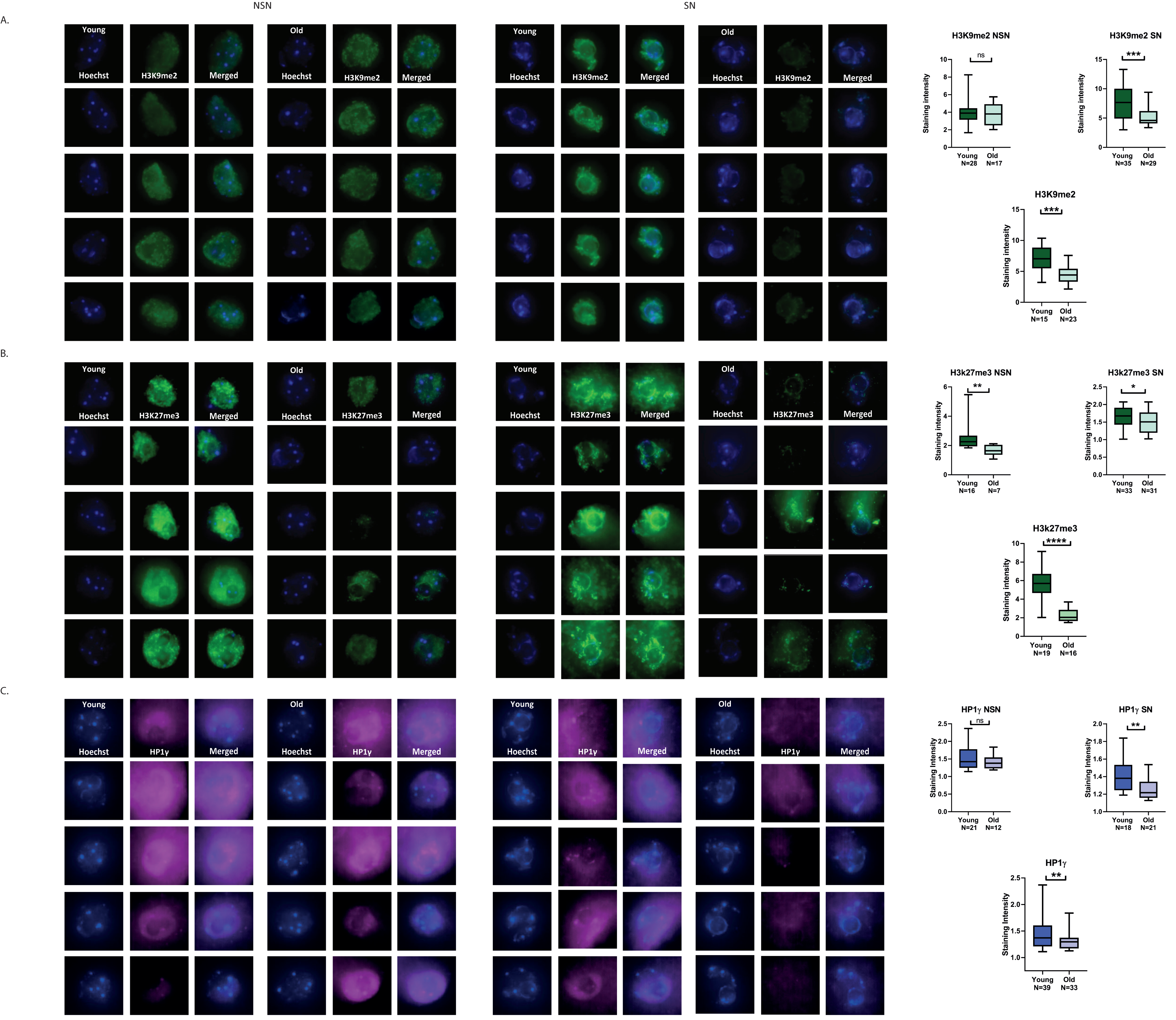


Figure S2: Comparison of heterochromatin staining in SN and NSN configured oocytes: (A) In situ staining of old and young GV mice oocytes for constitutive heterochromatin marker H3K9me2 in comparison between NSN and SN configuration of nuclei and staining quantification. (B) In situ staining of old and young GV mice oocytes for facultative heterochromatin marker H3K27me3 in comparison between NSN and SN configuration of nuclei and staining quantification. (C) In situ staining of old and young GV mice oocytes for constitutive heterochromatin marker HP1γ in comparison between NSN and SN configuration of nuclei and staining quantification.
