## Supplementary material for "Loss of heterochromatin and retrotransposon silencing constitute an early phase in oocyte aging": S1

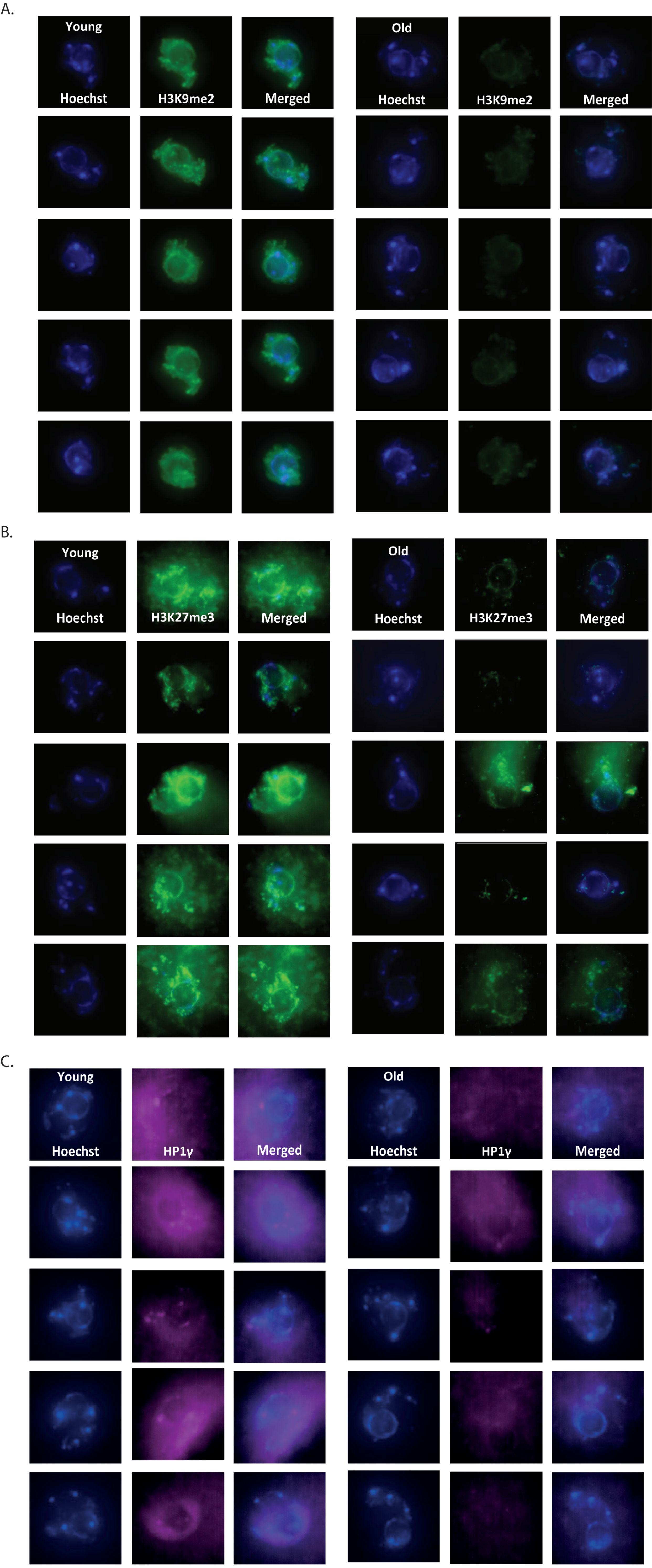


Figure S1: 9 months old oocytes lose heterochromatin marks (additional examples): (A) additional examples of in situ staining of old and young GV mice oocytes for constitutive heterochromatin marker H3K9me2. (B) additional examples of in situ staining of old and young GV mice oocytes for facultative heterochromatin marker H3K27me3. (C) additional examples of in situ staining of old and young GV mice oocytes for constitutive heterochromatin marker HP1γ.
